# DUSP11 is an intracellular innate immune checkpoint in lung adenocarcinoma

**DOI:** 10.1101/2025.01.24.634797

**Authors:** Brian J. Thomas, Xue Bai, Benjamin J. Cryer, Sydney M. Escobar, Lee-Ann H. Allen, Mark A. Daniels, Margaret J. Lange, Donald H. Burke

**Author notes:** Correspondence should be addressed to B.J.T or D.H.B. **Disclosures:** Nothing to disclose.

## Abstract

The discovery of immune checkpoints and the rapid growth of immuno-oncology (IO) have sparked tremendous efforts to utilize the immune system to treat a wide range of cancer types/subtypes. While the major focus of IO over the past decades has been to manipulate the adaptive immune system, recent attention has been given to manipulating the innate immune system to treat cancer and/or to enhance adaptive responses. In this work we detail the intracellular protein, Dual Specificity Phosphatase 11 (DUSP11), as an innate immune checkpoint (iIC) in Non-Small Cell Lung Cancer (NSCLC) adenocarcinoma (LUAD). Expression of this atypical phosphatase is correlated with patient survival for multiple cancer types, and we show here that its activity is important for the viability of lung cancer cells *in vitro*. Specifically, we demonstrate that DUSP11 knockdown in LUAD cells induces apoptosis and an innate immune response capable of activating other cells *in vitro,* and we provide evidence that these phenotypes are primarily mediated by the pattern recognition receptor, retinoic acid inducible gene I (RIG-I). Finally, we show that expression of DUSP11 is important for tumor engraftment and growth of human LUAD in mice. Overall, these data are the first to establish DUSP11 as an immunosuppressive, pro-neoplastic, and potentially targetable protein in LUAD. In addition, our data suggests that the anti-cancer mechanisms induced by diminishing the activity of DUSP11 are likely to be generalizable to other cancer types such as breast cancer, warranting future investigation and therapeutic development.

## Introduction

Over the past two decades, modern immunotherapies have revolutionized the way physicians and scientists think about treating cancer and have been the subject of a now rapidly developing area of study – immuno-oncology (IO). The major success of immune checkpoint inhibitors (ICIs) to treat cancer highlights (i) the efficacy of inhibiting immunosuppressive biological pathways as a means of re-activating the anti-tumor immune response and (ii) the generalizability of targeting such pathways, in that they are clinically useful across many cancer types. However, the successes of these therapies have not been without challenges. First, ICIs can induce potentially life-threatening side effects termed immune related adverse events (irAEs). Such adverse events are quite common with, on average, ∼20% of patients experiencing high grade irAEs and ∼40% experiencing any grade irAEs(1,2), although recently some irAEs have been associated with improved survival(3). Second, ∼40-70% of patients do not respond to ICIs(2), which is highly dependent upon cancer type. For example, microsatellite instable cancers have relatively high objective response rates(4), whereas oncogene (e.g., EGFR, AKT, KRAS) positive non-small cell lung cancer (NSCLC) rarely responds to immune checkpoint inhibitors, with most patients exhibiting progressive disease at follow up, including those previously treated with and responsive to targeted therapies but who developed resistance(5). Primary or secondary (acquired) resistance to ICIs is often attributed to the development of an immunologically cold tumor microenvironment (TME), suggesting that novel methods should be explored (i) to reactivate the immune system or (ii) to stimulate an inflammatory and anti-cancer TME, either alone or in combination with other therapeutics. Other adaptive immunity-based therapeutics have been developed to address some of these unmet needs, such as second and third generation T cell receptor (TCR) or chimeric antigen receptor (CAR) engineered cells, bispecific T cell engagers (BiTEs), and tumor infiltrating lymphocytes (TILs). However, these, too, have variable response rates, induce their own collection of irAEs, are typically patient specific (i.e., not generalizable), and aren’t yet especially good at treating solid tumors.

Every cell, including cancer cells, harnesses innate immune pathways which are typically thought of as serving anti-microbial roles. These pathways and their associated protein products are at the forefront of immunity, and they can promote the recruitment and activation of innate immune cells such macrophages, dendritic cells (DCs), and natural killer (NK) cells. In turn, these cells can play a role in either carrying out effector functions or in driving crosstalk with adaptive immune cells. Efficacy from targeting innate mediated immune pathways was first established in 2004 with the FDA approval of the Toll-like Receptor 7 (TLR7) agonist, Imiquimod, for the treatment of skin cancer. Since then, pre-clinical successes have been reported for various innate immune modulating strategies, and many clinical trials are underway to better understand the therapeutic potential of activating the innate immune system in cancer(6–10). Examples include using cytokines to promote APC activation/expansion, using targeted therapeutics to inhibit immunosuppressive proteins at the TME, and delivering agonists of pattern recognition receptors (PRRs)(7). Particularly for PRRs, oligonucleotides and oligonucleotide mimics have been developed as agonists for cGAS-STING(11), TLRs (12), and RIG-like receptors(13). However, these innate immune agonists must be calibrated to avoid extreme off-target effects and autoimmunity, as both of these factors can limit their therapeutic index (i.e., the ratio between the efficacious dose in cancer and the toxic dose in healthy cells). Evaluating alternative methods, such as targeting proteins that suppress innate immunity, is warranted.

Dual specificity phosphatase (DUSP11), also known as phosphatase interacting with RNA and ribonucleoprotein 1 (PIR1), is an atypical intracellular phosphatase with high affinity for immunostimulatory, 5’ triphosphate RNA (3pRNA)(14,15). DUSP11 is responsible for removing two phosphate groups from 3pRNA, a common agonist of the intracellular PRR, Retinoic acid Inducible Gene I (RIGI-I). The product of this reaction, 1pRNA, is relatively immunologically inert (16,17). DUSP11 has been extensively explored as a host protein that aids in virus escape from immune detection through this mechanism(16,18); however, this protein’s potential for mediating immune effects in cancer have largely escaped attention.

At least four lines of evidence point to a role for DUSP11 in cancer. (i) Expression of DUSP11 at the mRNA (https://www.cancer.gov/tcga) and protein(19) levels are correlated with patient survival data for multiple cancer types, such as LUAD, melanoma, renal, liver, head and neck, endometrial, pancreatic, and testicular cancers (**Figures 1A and S1**). (ii) Most cancers display a hallmark of dysregulated cellular energetics and metabolism(20) (e.g., altered 3pRNA transcript processing(21–23)). (iii) Anti-cancer and anti-viral immune mechanisms are significantly interconnected, and both can induce the effector functions of innate and adaptive immune cells for clearance. (iv) DUSP11 has recently been associated with pro-neoplastic processes in several different cancer types. With respect to this fourth point, DUSP11 was first described in relation to cancer in 2009 as a target molecule of p53, where its expression is upregulated upon exposure to DNA damaging agents and plays a role in p53-dependent inhibition of cell proliferation in osteosarcoma cells (24). In 2020, DUSP11 expression was shown to be upregulated in patient NSCLC tissue compared to healthy adjacent tissue, as well as in NSCLC cell lines compared to non-cancerous lung epithelial cells. This group showed that DUSP11 expression, which is correlated with NSCLC proliferation and migration, can be controlled directly by a microRNA (miR-513b-5p) that is complementary to *DUSP11* mRNA 3’UTR, or it can be controlled indirectly by a long non-coding RNA (lncRNA AZIN1-AS1) that acts on (i.e., suppresses) miR-513b-5p (25). In 2021, DUSP11 was shown to be an independent prognostic biomarker in patients with intrahepatic cholangiocarcinoma, although no mechanistic basis was elucidated(19). Finally, in 2024, DUSP11 was shown to be upregulated in advanced stage pancreatic ductal adenocarcinoma (PDAC) and contributed to gemcitabine (gem) resistance(26). The group reported that DUSP11 knockdown increased expression of vault RNAs, decreased phosphorylation of NFκB in response to gem treatment, and lowered protein kinase R (PKR) activity in the presence or absence of gem. As PKR-mediated NFκB phosphorylation has been proposed as a mechanism of gem resistance in PDAC, the group proposed a mechanism wherein the PKR-NFkB axis was perturbed by nc866, the expression of which is regulated by DUSP11 activity.

**Figure 1.**
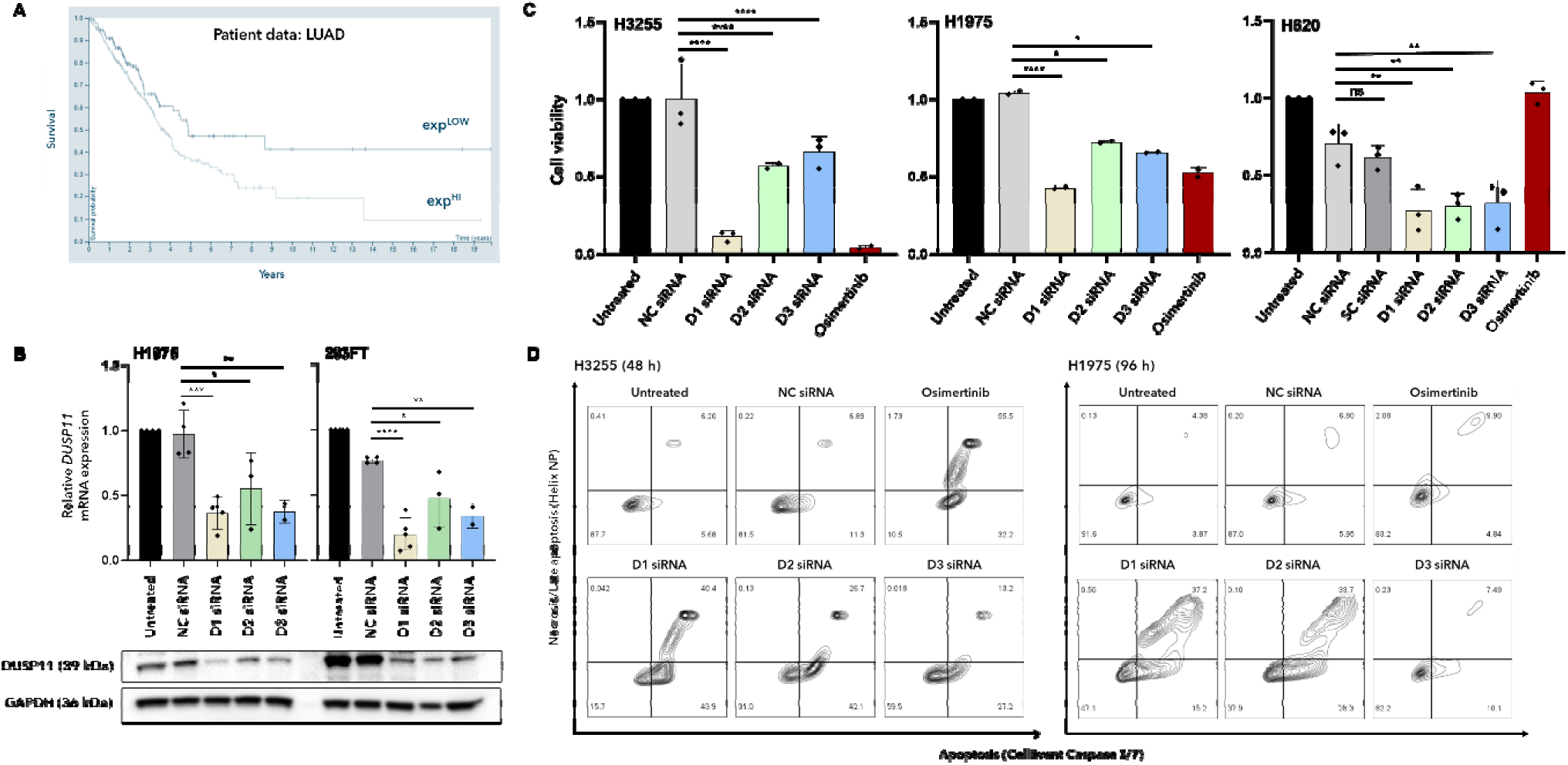
Knockdown of DUSP11 induces apoptosis in lung adenocarcinoma. (**A**) Kaplan-Meier survival curves for patents with lung adenocarcinoma (LUAD) having either high or low expression of *DUSP11* mRNA (Transcript Per Million, TPM cutoff = 19). Time (years) is represented on the *x*-axis and percent survival of the population is represented on the *y*-axis. The light blue line refers to high DUSP11 expression (above TPM cutoff) and dark blue lines refer to low DUSP11 expression (below TPM cutoff). Data were accessed from TCGA and obtained from the Human Protein Atlas. (**B**) LUAD (H1975) or non-cancerous embryonic kidney (293FT) cell line were left untreated (black bar) or treated with 10 nM non-targeting control (NC; grey bar) or various Dicer substrate siRNA targeting DUSP11 mRNA (D1, D2, D3 siRNA; sand, green, blue bars, respectively), then mRNA (24 h) and protein (48 h) were assessed using RT-qPCR or western blot, respectively. For mRNA, all values are normalized to GAPDH and reported relative to mRNA from the untreated sample on the *y*-axis. Plotted values represent mean ± SD for n=2-4 independent experiments, each with 2-3 technical replicates. D1, D2, and D3 siRNA but not NC siRNA decreased both mRNA and protein expression of DUSP11. (**C**) LUAD (H1975, H3255, H820) cell lines were treated with 10 nM NC (light grey bar), scramble (SC; dark grey bar), D1 (sand bar), D2 (green bar), D3 (blue bar) siRNA or 0.5 µM Osimertinib (red bar) for 96 h (media change with no siRNA or drug at 48 h), and cell viability was assessed by MTS assay. Cell viability relative to untreated cells is reported on the *y*-axis; plotted values represent mean ± SD for n=2-3 independent experiments, each with 2-4 technical replicates. Cells treated with D1, D2, and D3 siRNA exhibited decreased cell viability compared to controls. (**D**) LUAD cells were treated with 20nM NC, D1, D2, or D3 siRNA for 48 h (H3255) or 96 h (H1975) or 0.1 µM Osimertinib for 48 h (both cell lines) and assessed by flow cytometry for CellEvent™ Caspase 3/7 reagent staining (apoptosis; *x*-axis) or Helix NP™ staining (cell impermeable dye for necrosis/late apoptosis; *y*-axis). Representative contour plots in which percent of cells unstained (Q3) or stained with Caspase 3/7 reagent only (Q4), Helix NP™ only (Q2), or both reagents (Q1) from n=2 independent experiments for each cell line are shown. Cells treated with D1, D2, and D3 siRNA increased percent of cells undergoing apoptosis compared to controls. D1 siRNA was deemed the most efficacious reagent when evaluating cell viability. For B-C, statistical analysis was performed using a one-way ANOVA with post hoc multiple comparisons test corrected using the Dunnett method (*p < 0.05, **p < 0.01, ***p < 0.001, ****p < 0.0001).

While these recent studies highlight an associated role for DUSP11 in cancer, its role in modulating the innate immune system has not yet been described in the context of cancer as it has so readily been done with viral infection. We hypothesized that DUSP11 knockdown would increase the presence of immunostimulatory RNA such as 3pRNA and that detection of these RNAs by PRRs would activate anti-cancer immune mechanisms. Our findings lead us to describe DUSP11 as a targetable, innate immune checkpoint (iIC) in non-small cell lung cancer (NSCLC) adenocarcinoma (LUAD), and likely for other cancers. LUAD is a leading cause of death worldwide, largely due to mutant oncogene acquired therapeutic resistance(27). Lack of effective alternative therapeutic options, including ICIs(5,28), presents a clear unmet need for novel methods to treat refractory LUAD. We show that multiple Dicer substrate small interfering RNAs targeting DUSP11 effectively decreased expression at the mRNA and protein levels and that these treatments had significant impacts on the cell viability of at least two cancer cell types. In LUAD, this decrease in cell viability was due to induction of apoptosis, and innate immune responses were significantly upregulated, as evidenced by: (i) release of type I interferon (IFN), (ii) increased expression of interferon-stimulated genes (ISGs), and (iii) paracrine stimulation both of cells expressing an interferon-stimulated gene response element (ISRE) reporter and of human peripheral blood mononuclear cells (hPBMCs) *in vitro*. As DUSP11 knockdown has been shown to upregulate 3pRNA in cancer(26), we suspected that the PRR, RIG-I, was responsible for mediating the anti-cancer phenotype, similar to its role in anti-viral responses(16). Indeed, we show that the knockdown of RIG-I, but not other intracellular PRRs, recovered the cell viability and innate immune response phenotypes. Corroborating this finding, we show that knocking down DUSP11 sensitized cells to the cytotoxic effect of RIG-I agonizing 3pRNA. Furthermore, LUAD cells engineered to be deficient in RIG-I using CRISPR/Cas9 did not respond to DUSP11 knockdown. Finally, we demonstrate that knockdown of DUSP11 prior to LUAD cell implantation in mice significantly decreased tumor engraftment and tumor burden. Overall, our work establishes DUSP11 as an excellent target for the immunomodulation of LUAD and suggests that the innate immune mechanisms induced by modifying DUSP11 expression may be shared across multiple cancer types.

## Results

### Expression of DUSP11 is correlated with patient survival for multiple cancer types

In alignment with our hypothesis stated above, we speculated that patients with higher expression of DUSP11 would display poorer survival compared to those with lower expression for some cancer types, as DUSP11 might support an immunosuppressive or immunologically cold phenotype within their tumors. As detailed above, this speculation was supported by previous studies that examined DUSP11 expression as a prognostic biomarker in a subset of liver(19) or pancreatic cancers(26). Utilizing data generated by The Cancer Genome Atlas (TCGA) Research Network (https://www.cancer.gov/tcga) and obtained from the Human Protein Atlas (https://www.proteinatlas.org), we identified multiple cancer types for which lower DUSP11 expression correlated with improved survival, regardless of stage (**Figure 1A and S1A-G**). These cancers included LUAD (**Figure 1A**), melanoma, renal (mainly chromophobe and papillary), liver, head and neck, endometrial, pancreatic, and testicular cancers (**Figure S1A-G**). There were also cancer (sub)types that did not correlate, or that had the opposite correlation (**Figure S1H-J**). These included lung squamous cell carcinoma (LUSC, **Figure S1H**; LUSC is a form of NSCLC that is pathologically different from LUAD), breast cancer (**Figure S1I**), and colorectal cancer (CRC, mainly rectal; **Figure S1J**). Nevertheless, these TCGA data suggest that DUSP11 expression is correlated with patient survival for multiple cancer types and that an underlying mechanism involving this protein may play a role.

### DUSP11 knockdown decreases cell viability of lung adenocarcinoma

To establish whether the expression of DUSP11 was important for LUAD cell viability, we first evaluated three Dicer substrate small interfering RNAs (siRNAs) targeting various loci on *DUSP11* mRNA (D1, D2, D3 siRNA) for their capacity to knock down DUSP11 in either HEK293FT cells (non-cancerous embryonic kidney cell) or H1975 cells (LUAD cell). We chose to develop these as Dicer substrate siRNAs rather than conventional 21mer siRNAs as this approach has been demonstrated to improve RISC loading and potency of target knockdown(29,30), and they can readily be integrated into targeted delivery reagents(31). Compared to non-targeting control (NC) siRNA, a single treatment (transfection) of 10nM D1, D2, or D3 siRNA was effective at knocking down DUSP11 at the mRNA (24 h) and protein (48 h; **Figure 1B**) levels by ∼50-80%, depending upon the targeted sites within the *DUSP11* gene.

We next evaluated the impact of DUSP11 knockdown on cell viability of various LUAD cancer cell lines (H1975, H3255, H820, A549) using MTS assays (**Figure 1C and S2A**). To ensure observed effects were due to gene silencing rather than non-specific or off-target effects of RNA transfection, all DUSP11 targeting siRNAs were compared to either NC siRNA or a scrambled control (SC) siRNA with the same nucleotide composition of D1 siRNA but with a shuffled sequence. Compared to these controls, a single treatment of 10nM D1, D2, or D3 siRNA decreased cell viability of three of the four LUAD cell lies (H1975, H3255, H820; **Figure 1C**) when assessed at 96 h. No difference in cell viability was observed for A549 (**Figure S1A**). D1 siRNA was the most efficacious reagent of the three siRNAs, decreasing cell viability by ∼60-90% (∼1-5-fold decrease relative to D2 or D3 siRNA), depending on the cell line.

As MTS based cell viability assays only determine the number of metabolically active cells, we were interested in understanding whether cells treated with D1 siRNA were undergoing apoptosis or another cell death mechanism. Therefore, we treated two LUAD cells lines – H3255 and H1975 – with 20nM D1 siRNA for 48 h or 96 h, respectively, and analyzed the proportion of cells stained for mature caspase 3/7 (Cell Event™; apoptosis) and/or or a cell impermeable dye (Helix NP™; necrosis/late apoptosis). Compared to non-targeting controls, all siRNAs targeting *DUSP11* mRNA increased the number of cells containing mature caspase 3/7 ± Helix NP™, similar to cells treated with Osimertinib (**Figure 1D**), which is known to induce apoptosis(32). These results suggest that DUSP11 knockdown induces apoptosis in LUAD and that its expression plays an important role in cell survival. Consistent with MTS cell viability assays, D1 siRNA was the most efficacious reagent compared to D2 or D3 siRNA.

### DUSP11 knockdown decreases tumor engraftment and tumor burden in mice harboring lung adenocarcinoma xenografts

We next sought to establish whether the anti-cancer phenotype observed *in vitro* upon DUSP11 knockdown was reproducible within a more biologically relevant system, or if additional variables, such as the production of an immunosuppressive TME, might counteract these effects. To this end, H1975 cells were treated with a single 10 nM dose of NC siRNA, D1 siRNA, or left untreated for 24 h, and ∼2.5M live cells (Cell Event™ caspase 3/7 and Helix NP™ negative) were implanted into the right flank of BALB/c nude mice. Tumor growth kinetics and mouse weight were then tracked until at least one mouse reached a humane endpoint (**Figure 2A**). Average tumor volumes (**Figure 2B**) of individual tumor trajectories (**Figure 2C and S4A**) revealed that mice implanted with cells treated with D1 siRNA had significantly lower tumor burden compared to mice implanted with untreated cells (*p* < 0.0001) or NC siRNA treated cells (*p* < 0.0001) when assessed by a simple linear regression model. An area under the curve (AUC) model also revealed a significant tumor burden reduction in response to DUSP11 knockdown relative to mice implanted with untreated cells (*p* = 0.0494) or NC siRNA treated cells (*p* = 0.0085) (**Figure S4B**). At the humane endpoint (day 30 post injection), all mice were euthanized and imaged in prone and right recumbent anatomical positions (**Figure S4D**), then tumors were excised (**Figure 2D**) and ex vivo tumor weights were obtained (**Figure S4C**). Mice implanted with cells treated with D1 siRNA (mean = 0.066 g) had significantly lower *ex vivo* tumor weights when compared to mice implanted with untreated cells (mean = 0.448 g, *p* = 0.035) but not NC siRNA treated cells (mean = 0.234 g, *p* = 0.113). Together, these data support that DUSP11 expression significantly influences tumor engraftment and/or growth.

**Figure 2.**
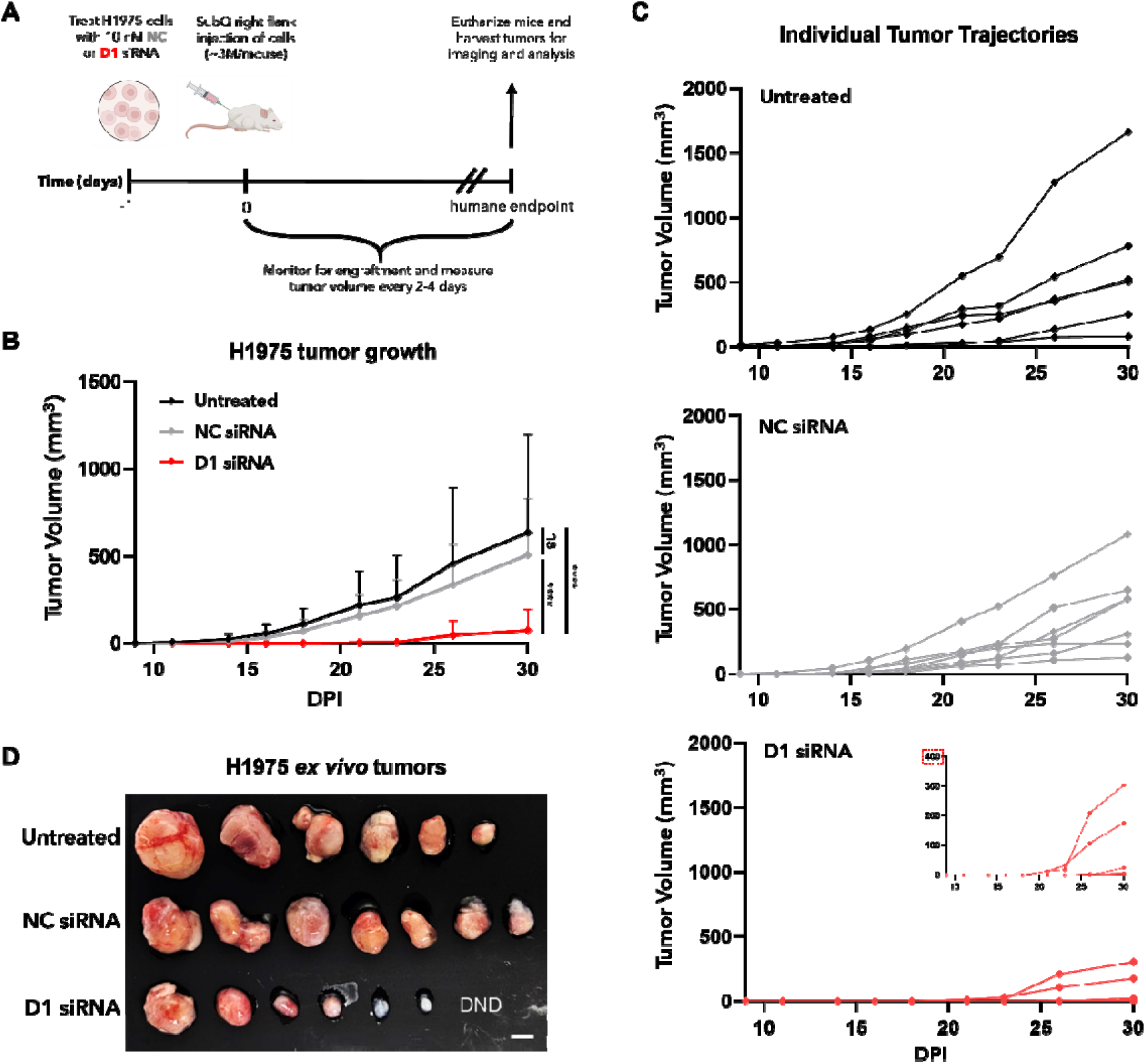
DUSP11 is important for tumor cell engraftment and growth in mice implanted with lung adenocarcinoma. (**A**) Schematic of mouse model assessing LUAD cell engraftment and growth in BALB/c nude mice. In brief, mice were implanted with ∼ 2.5M live H1975 cells that were either left untreated or treated with 10nM NC or D1 siRNA for 24 h. Live cells were determined by negative staining for Helix NP™ and CellEvent™ Caspase 3/7 reagent. Tumor engraftment and mouse weight were then tracked until a humane endpoint (day 30). (**B-C**) Average (B) or individual trajectories (C) of tumor burden of mice implanted with H1975 cells left untreated (black lines) or treated with 10 nM NC (grey lines) or D1 (red lines) siRNA for 24 h. (**D**) *Ex vivo* image of H1975 tumors from all mice at the endpoint (DND = tumor did not develop). For B-C, days post implantation (DPI) is reported on the *x*-axis and tumor volume (mm^3^ = 0.5 x L x W^2^) is reported on the *y*-axis. For B, plotted values represent mean ± SD for n=6 (untreated) or 7 (NC and D1 siRNA treated) mice from each treatment group. Mice implanted with cells treated with D1 siRNA exhibited a significantly lower tumor burden compared to those implanted with untreated cells or cells treated with NC siRNA. Mouse weights remained relatively consistent throughout the experiment (data not shown). Statistical analysis was preformed using a linear regression model (****p < 0.0001).

BALB/c nude mice are only partially immunodeficient, lacking adaptive immunity (T cells) but still harboring a relatively intact innate immune system. Therefore, it is plausible that immune cells interacting with the implanted cancer cells could have impacted tumor engraftment and/or growth. As our experimental model prevented direct assessment of tumor infiltrating immune cells, we sought to assess the activation status of circulating immune cells in the blood at 24 h and 7 days post cancer cell implantation. Changes in the relative percentages of circulating B cells, monocytes, DCs, and NK cells, along with their activation states, as determined by MFI of CD69, CD86, and MHC class II, were assessed by flow cytometry (**Figure S5**). There were no significant differences in relative cell populations (**Figure S5A**) or the cell surface levels of CD69 or CD86 for any circulating cell type at either timepoint (**Figure S5B-C**). Interestingly MHC class II levels were significantly increased, albeit variably, on monocytes, B cells, and NK cells in mice implanted with siRNA treated cells compared to those implanted with untreated cells; however, only the increase in NK cells was specific to treatment with D1 siRNA (**Figure S5B-C**). NK cells constitute an innate immune cell population that is more sensitive to type I IFN stimulation than are the other cell types; however, NK cell activation by Type I IFN is typically characterized by increased proliferation and effector function (e.g., upregulation of CD69)(33,34); unfortunately, increased MHC class II expression on NK cells has not been well characterized. As confounders to precise interpretation, many of the cytokines that are produced from human tumors have variable cross reactivity with murine immune cells and receptors(35), and the nude mouse innate immune system is still not as robust as an immunocompetent mouse. Therefore, while the small differences noted in activation of circulating immune cells are not unexpected, these confounders make it difficult to interpret precisely and require further evaluation, ideally by evaluating tumor infiltration by immune cells in an immunocompetent, humanized mouse model and where tumors are treated *in situ* with a DUSP11 targeting reagent.

### Knockdown of DUSP11 induces an innate immune response in lung adenocarcinoma

As DUSP11 expression and activity have been previously reported to suppress anti-viral immune responses, we were interested in determining whether DUSP11 expression also played a similar role in suppressing anti-cancer immune responses. A key phenotype of an innate immune response is the production of type I IFNs(34,36). Therefore, we first evaluated the production of IFNβ by H3255 and H1975 cells treated with D1 siRNA. Transfecting cells with 10nM D1 siRNA significantly increased the amount of IFNβ released into the media at 48 h relative to controls (**Figure 3A**; H3255: 3.9 ng/ml, p < 0.0001; H1975 2 ng/ml, p < 0.0001). Type I IFNs have been shown to enhance both innate and adaptive anti-cancer immune responses, both in treatment naïve cancers and in overcoming therapeutic resistance (34,37–40); therefore, we were next interested in understanding whether the cytokines produced as a result of knocking down DUSP11 (such as IFNβ) could activate other cells in a paracrine fashion. We transfected HEK293FT cells with an ISRE dual-luciferase reporter plasmid and exposed them for 16 h to conditioned media taken from siRNA-treated LUAD cells. Media from LUAD cells transfected with 10 or 20nM D1 siRNA significantly increased luciferase production in HEK293FTs (**Figure 3B and S6A-B;** H3255: ∼7-fold increase over control siRNA; H1975: ∼6-fold increase). To assure that this observation was not due to changes in HEK239FT DUSP11 expression from residual D1 siRNA in the media, we assessed luciferase production by HEK293FTs incubated with the same media from H1975 cells but treated with RNase (to remove any residual siRNA) or unconditioned media with 10 nM freshly complexed siRNA (to show effect of knocking down DUSP11 in HEK293FT on luciferase production; **Figure S6B**). As expected, treatment of conditioned media with RNase had nearly no effect on luciferase production when compared to conditioned media not treated with RNase (still a ∼5-fold increase), whereas treatment with fresh D1 siRNA did not stimulate significant luciferase production. Both of these controls support the interpretation that the effects observed above using conditioned media were driven by cytokines in the media rather than by alteration of DUSP11 expression in the reporter cells. Overall, these data support that cytokines/chemokines produced by DUSP11 knockdown in LUAD are sufficient for paracrine stimulation and could play a role in crosstalk with other immune cells, activating the anti-cancer innate and adaptive immune systems.

**Figure 3.**
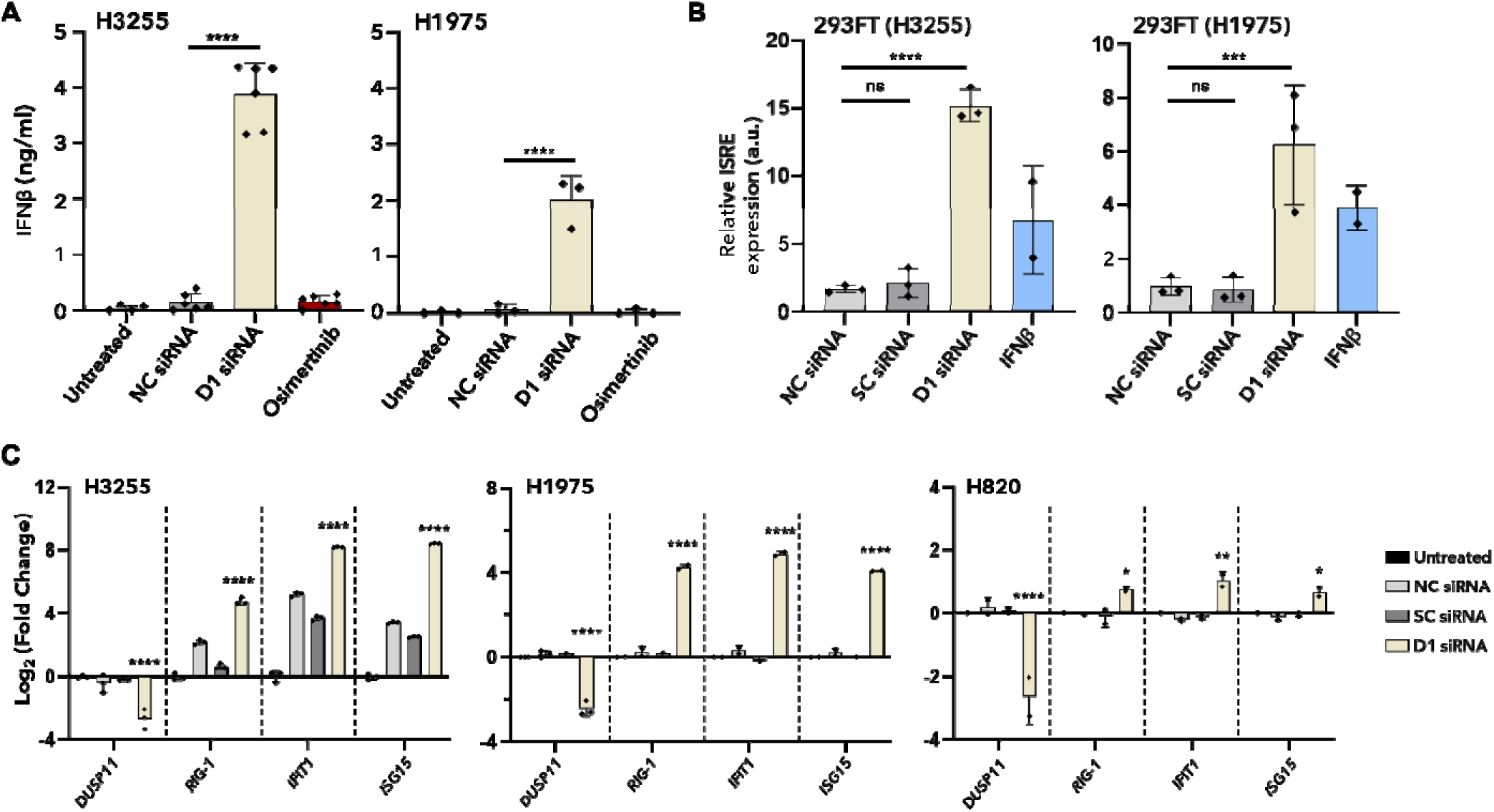
DUSP11 knockdown induces an innate immune response in lung adenocarcinoma. (**A**) LUAD cells lines (H3255 and H1975) were left untreated (black bar) or treated with either 20 nM NC (grey bar) or D1 (sand bar) siRNA, or 0.5 uM Osimertinib (red bar) for 48 h. Production of and release of IFNβ into the media was assessed by enzyme linked immunosorbent assay (ELISA). IFNβ concentration (ng/mL) as determined by a standard curve using recombinant IFNβ is reported on the *y*-axis and plotted values represent mean ± SD for n=3-6 independent experiments, each with 2-4 technical replicates. Cells treated with D1 siRNA produced significantly more IFNβ compared to those left untreated or those treated with NC siRNA or Osimertinib. (**B**) HEK293FT transfected with an ISRE reporter were treated with media from LUAD cells transfected with either 10 nM (H3255) or 20 nM (H1975) NC (light grey bar), SC (dark grey bar), D1 (sand bar) siRNA, or 100 ng/mL IFNβ for 48 h. IFN-stimulated transcription from the ISRE was determined by monitoring luciferase substrate conversion (light production) and is reported relative to untreated cells on the *y*-axis (a.u. = arbitrary units). Plotted values represent mean ± SD for n=2-3 independent experiments, each with 2-3 technical replicates. Media from D1 siRNA treated cells exhibited significantly increased reporter expression compared to media from control treated cells. (**C**) LUAD cells were left untreated (black bar) or treated with either 10 nM (H3255, H820) or 20 nM (H1975) NC (light grey bar), SC (dark grey bar), or D1 (sand bar) siRNA for 24 h and mRNA expression was assessed using RT-qPCR. All values are normalized to GAPDH and reported relative to mRNA from the untreated sample (Log_2_(Fold Change = ΔΔCq)) on the *y*-axis. Plotted values represent the mean ± SD for n=2-3 independent experiments, each with 2-3 technical replicates. Cells treated with D1 siRNA exhibited a significant increase in expression of ISGs compared to control siRNAs. For A-B, statistical analyses were performed using a one-way ANOVA with post hoc multiple comparisons test corrected using the Dunnett method (***p < 0.001, ****p < 0.0001). For C, statistical analysis was performed using a two-way ANOVA with post hoc multiple comparisons test corrected using the Dunnett method (*p < 0.05, **p < 0.01, ****p < 0.0001) and are reported relative to NC siRNA.

Another hallmark of an innate immune response is the increased expression of interferon-stimulated genes (ISGs)(41,42). DUSP11 expression has been previously reported to be important for transcription of ISGs encoding IFIT1 (*IFIT1*) and ISG15 (*ISG15*) in response to viral infection (16,43). Therefore, we assessed whether the expression of these ISGs, as well as the gene encoding RIG-I (*DDX58*), was affected in response to DUSP11 knockdown in various LUAD cell lines (H1975, H3255, H820). Across all cell lines, knocking down DUSP11 significantly increased the expression of *DDX58, IFIT1,* and *ISG15* when compared to NC or SC siRNA (**Figure 3C**); however, in H3255, transfection of controls also increased expression of these target genes, albeit to a lesser extent. As the cell viability phenotype in H3255 was only present upon D1 siRNA treatment and not control siRNA treatment, this suggests that while changes in ISG expression are related to DUSP11 expression, they may only be only partially responsible for the observed anti-cancer effect, and that other elements such as changes at the protein level (i.e., phosphorylation, ubiquitination) may also be involved. Nevertheless, taken together these data support that LUAD cells in which DUSP11 has been knocked down experience a robust innate immune response, as evidenced by upregulation of ISGs and the production of type I IFN, analogous to the robust innate immune responses observed in virally infected cells.

### RIG-I is required for the innate immune response and cell viability phenotype

The same 3pRNA that serves as a substrate of DUSP11(15,16) is also the immunostimulatory ligand of RIG-I(44,45). While exogenous 3pRNA can accumulate during infection by certain viruses, RNA polymerase III (RNAPIII) is the primary protein responsible for producing most host-derived, endogenous 3pRNA pools(46). In cancer, dysregulated transcription by RNAPIII has been described(15,21,22), and its increased expression is correlated with poorer survival in some cancers(47) (TCGA). Recently, vault RNAs (vtRNAs) transcribed by RNAPIII (48–50) have been associated with DUSP11 in pancreatic cancer (26) and in crosstalk with innate immune receptors, specifically RIG-I (43,51). Given these known interactions, it is noteworthy that RIG-I expression is increased upon knockdown of DUSP11 in our LUAD cell lines (*see previous section*). To test the possibility that RIG-I was mediating the innate immune response and cell viability phenotype observed above, LUAD cells (H3255, H1975, or H820) were treated with 10 nM NC or D1 siRNA in the presence or absence of 10 nM RIG-I Dicer substrate siRNA (RIG-I siRNA), and we evaluated the impact of this double knockdown on cell viability, expression of ISGs, and production of type I IFN. These studies revealed that the decrease in cell viability (96 h; **Figure 4A-B**), increase in ISG expression (48 h; **Figure 4C**), and release of IFNβ (48h; **Figure 4D**) observed upon DUSP11 knockdown were all partially recovered when RIG-I was also knocked down (**Figure 4C and S7A**). RIG-I knockdown alone did not have significant effects on cell viability, ISG expression, or IFNβ production. In contrast to RIG-I, double knockdown using dicer substrate siRNAs targeting either *IFIH1* mRNA (melanoma differentiation-associated protein 5, ‘MDA5 siRNA’) or *EIF2AK2* mRNA (Protein kinase R, ‘PKR siRNA’) had no effect on the anti-cancer phenotype induced by DUSP11 knockdown at 48 h or 96 h in H3255 and H1975 cells (**Figure S7B**), suggesting that these RNA stimulated intracellular PRRs(41,52,53) were not appreciably involved in the mechanisms governing LUAD cell viability. During anti-viral signaling, the RIG-I N-terminal tandem caspase activation recruitment domains (CARDs) interact with mitochondrial antiviral signaling protein (MAVS)(54). Unfortunately, knocking down MAVS had significant impacts on LUAD cell viability (**Figure S7B**), precluding straightforward interpretation of its role in mediating signaling through the DUSP11/3pRNA/RIG-I axis. Nevertheless, the data above support the notion that RIG-I mediates both the anti-cancer phenotype and innate immune response upon DUSP11 knockdown.

**Figure 4.**
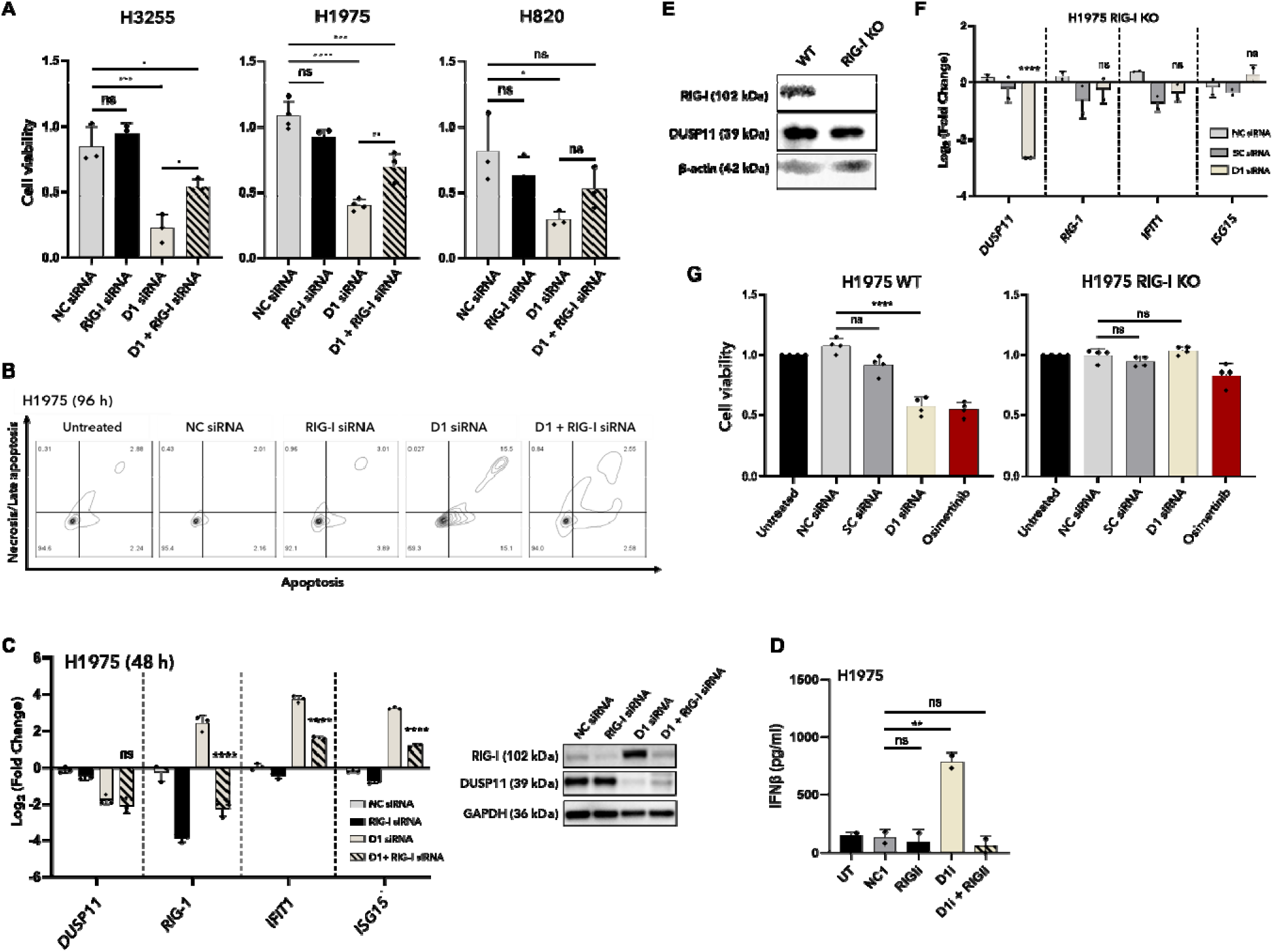
RIG-I is required for the innate immune response and cell viability phenotype induced upon DUSP11 knockdown. (**A**) H3255, H1975, and H820 cells were transfected with 10 nM NC (grey bar) or D1 (sand bar) siRNA in the presence or absence of 10nM RIG-I siRNA (black or black/sand bar) for 96 h (media change with no siRNA at 48 h), and cell viability was assessed by MTS assay. Cell viability relative to untreated cells is reported on the *y*-axis and plotted values represent mean ± SD for n=3-4 independent experiments, each with 2-4 technical replicates. (**B**) H1975 cells were treated as in A for 96 h and assessed by flow cytometry for CellEvent™ Caspase 3/7 reagent staining (apoptosis; *x*-axis) or Helix NP™ staining (cell impermeable dye for necrosis/late apoptosis; *y*-axis). Representative contour plots in which percent of cells unstained (Q3) or stained with Caspase 3/7 reagent only (Q4), Helix NP™ only (Q2), or both reagents (Q1) from n=3 independent experiments for each cell line are shown. (**C**) H1975 cells were treated as in A for 48 h, and mRNA (*left*) and protein (*right*) expression were assessed using RT-qPCR and western blot, respectively. For mRNA, all values are normalized to GAPDH and reported relative to mRNA from the untreated sample (Log_2_(Fold Change = ΔΔCq)) on the *y*-axis. Plotted values represent the mean ± SD for n=3 independent experiments, each with 2-3 technical replicates. For protein, representative blots from n=2 independent experiments are shown. Target proteins were knocked down as expected relative to the loading control, GAPDH. (**D**) H1975 cells were treated as in A for 48 h and the production and release of IFNβ into the media was assessed by enzyme linked immunosorbent assay (ELISA). IFNβ concentration (ng/mL) as determined by a standard curve using recombinant IFNβ is reported on the *y*-axis. Plotted values represent mean ± SD for n=2 independent experiments, each with 4 technical replicates. For A-D, the decrease in cell viability (A), increase in apoptosis (B), increase in ISG expression (C), and increase in IFNβ release (D) upon treatment with D1 siRNA was partially recovered when cells were co-transfected with RIG-I siRNA. (**E**) H1975 cells were engineered to be knocked out for RIG-I using CRISPR/Cas9 (*see Figure S8 for additional details*). Untreated parental (WT) and RIG-I deficient (RIG-I KO) H1975 cells were lysed, and protein expression of RIG-I and DUSP11 was assessed by western blot. Representative blots from n=2 independent experiments are shown. RIG-I was undetectable in RIG-I KO cells relative to WT (parental) cells and the loading control, β-actin. DUSP11 expression was slightly decreased in RIG-I KO cells. (E-F) H1975 RIG-I KO cells were transfected with 10 nM NC, SC or D1 siRNA and mRNA expression (24 h; E) and cell viability (96 h with 48 h media change; F) were assessed as above. Unlike parental cells, H1975 RIG-I KO cells did not exhibit increased ISG expression or decreased cell viability when transfected with D1 siRNA when compared to controls. For A, D, and G, statistical analyses were performed using a one-way ANOVA with post hoc multiple comparisons test corrected using the Šidák (A) or Dunnett (D,G) method (*p < 0.05, **p < 0.01, ***p < 0.001, ****p < 0.0001). For C, statistical analysis was performed using a two-way ANOVA with post hoc multiple comparisons test corrected using the Dunnett method (****p < 0.0001) and are reported relative to D1 siRNA.

To further establish reliance of the observed anti-cancer phenotype and innate immune responses on RIG-I, and to decipher whether the partial recovery phenotype noted above was due to incomplete knockdown, we engineered RIG-I knockout (RIG-I KO) H1975 cells (**Figure 4E and S8A-B**), then treated these cells with either 10nM NC or D1 siRNA and assessed both ISG expression (**Figure 4F**) and cell viability (**Figure 4G**). In agreement with double knockdown studies and in contrast to observations in H1975 WT (parental) cells, the knockdown of DUSP11 did not significantly influence the expression of *ISG15* or *IFIT1* from H1975 RIG-I KO cells at 48 h (**Figure 4F**) and had no significant effect on the cell viability of these cells at 96 h (**Figure 4G**), when compared to control siRNAs. These data provide strong evidence that in LUAD, DUSP11 modulates both RIG-I expression and activity, which in turn mediates the anti-cancer phenotype and innate immune responses, likely as a result of shifting balances in immunostimulatory 3pRNA pools. Considering our findings in a RIG-I KO background and that RIG-I mRNA was only partially knocked down after 24 h co-transfection in H1975 (**Figure S7C**), we conclude that the incomplete recovery of the DUSP11 knockdown phenotype observed above when co-transfected with RIG-I siRNA was due to inadequate RIG-I knockdown.

### DUSP11 expression controls sensitivity of lung adenocarcinoma cells to immunostimulatory 5’ triphosphate RNA

Recent works have shown that exploiting RLR pathways is a feasible method to activate anti-cancer immune responses(13). While radiation and chemotherapy have been described to activate these pathways, much of the recent work in this field has focused on the development of methods and/or reagents to deliver RLR stimulating ligands/agonists (e.g., 3pRNA) including the use of intratumoral injection, nanoparticles, and even CAR T cells. As DUSP11 knockdown likely increases the pool of available immunostimulatory 3pRNA, we considered that even minor changes in DUSP11 expression could increase cancer cell sensitivity to RIG-I agonizing 3pRNA. To test directly the impact of manipulating 3pRNA, we first established a dose of D1 siRNA that would be tolerated by two LUAD cell lines (H3255 and H820). Transfecting 1 nM D1 siRNA had little effect on the viability of these cells, while transfecting 10 nM D1 siRNA reduced cell viability too much to allow monitoring of the effects of 3pRNA (data not shown). We then co-transfected these two LUAD cell lines with either 1 nM NC siRNA or D1 siRNA and various doses of a commercial 3pRNA RIG-I agonist (InvivoGen 3p-hpRNA) and then assessed cell viability by MTS assay at 48 h (**Figure 5**). In both cell lines, the agonist was cytotoxic with EC50 ∼140 µg/mL. However, suboptimal DUSP11 knockdown using 1 nM D1 siRNA, but not NC siRNA, decreased EC50 for the agonist nearly 2-fold to ∼80 µg/mL (**Figure 5**). These data suggest that modulating 3pRNA levels by targeting DUSP11 expression is a feasible way to sensitize cancer cells to RIG-I agonists.

**Figure 5.**
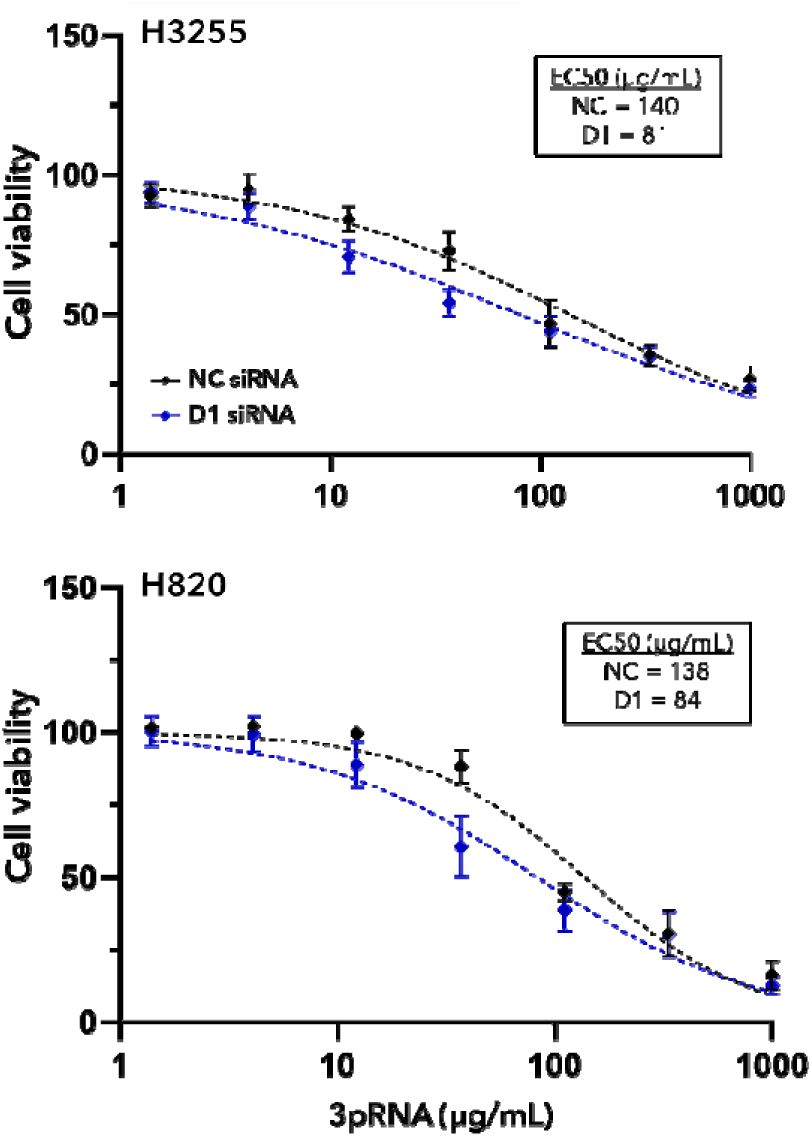
DUSP11 knockdown sensitizes cells to exogenous 3pRNA. H3255 (*top*) or H820 (*bottom*) cells were co-transfected with 1nM NC (black lines) or D1 (blue lines) siRNA and a series of doses of 3pRNA (RIG-I agonist) and cell viability was assessed by MTS assay. Cell viability relative to untreated cells is reported on the *y*-axis. Plotted values represent mean ± SD for n=2-3 independent experiments, each with 2-4 technical replicates. EC50 values were determined by fitting a non-linear curve using a least squares regression method. LUAD cell lines treated with D1 siRNA and 3pRNA exhibited a ∼2-fold decrease in EC50 values when compared to those treated with NC siRNA. Statistical analyses were performed using an extra sum of squares F-test on the fit curves (H3255: p = 0.0238; H820: p = 0.0174).

### Cytokines produced by lung adenocarcinoma cells upon DUSP11 knockdown can activate adaptive immune cells

As shown above, media from D1 siRNA treated LUAD cells significantly increased luciferase expression in a HEK293FT-ISRE reporter system (**Figure 3B**). To test whether similar immunomodulatory paracrine effects occur in a more clinically translatable model, we assessed whether media from these cells could activate hPBMCs (**Figure 6**). H1975 cells were treated with 10 nM NC or D1 siRNA in the presence or absence of 10 nM RIG-I siRNA, then diluted cell-free media from these cultures was incubated with hPBMCs from two donors for ∼20 h and adaptive immune cell states and activation status were assessed. For both donors, both B and T cell activation (CD69 positivity) was upregulated upon incubation with media from D1 siRNA treated cells. In contrast, little (donor 2) or no (donor 1) activation was observed with media from NC siRNA transfected cells or from D1 and RIG-I siRNA co-transfected cells (**Figure 6A-B**). Activation of both memory and naïve T cells was upregulated, with CD8 memory cells showing the highest percentage of CD69 positivity (**Figure 6B**). While minimal changes in the percentage of T central memory (T_CM_) and T effector memory (T_EM_) cells were noted in PBMCs from donor 2, PBMCs from donor 1 showed a small increase in both CD4 and CD8 T_CM_ populations. These findings suggest that cytokines produced by LUAD cells upon DUSP11 knockdown, but not those produced in the presence of RIG-I knockdown, can activate adaptive immune cells in paracrine fashion and may alter their lineage differentiation (i.e., T_CM_ versus T_EM_). As hPBMCs consist of various cell types (B and T cells, monocytes, DCs, NK cells) and only the adaptive immune cells were assessed here, characterizing the changes of and interactions among all cell types will be of interest for future studies, especially *in vivo*.

**Figure 6.**
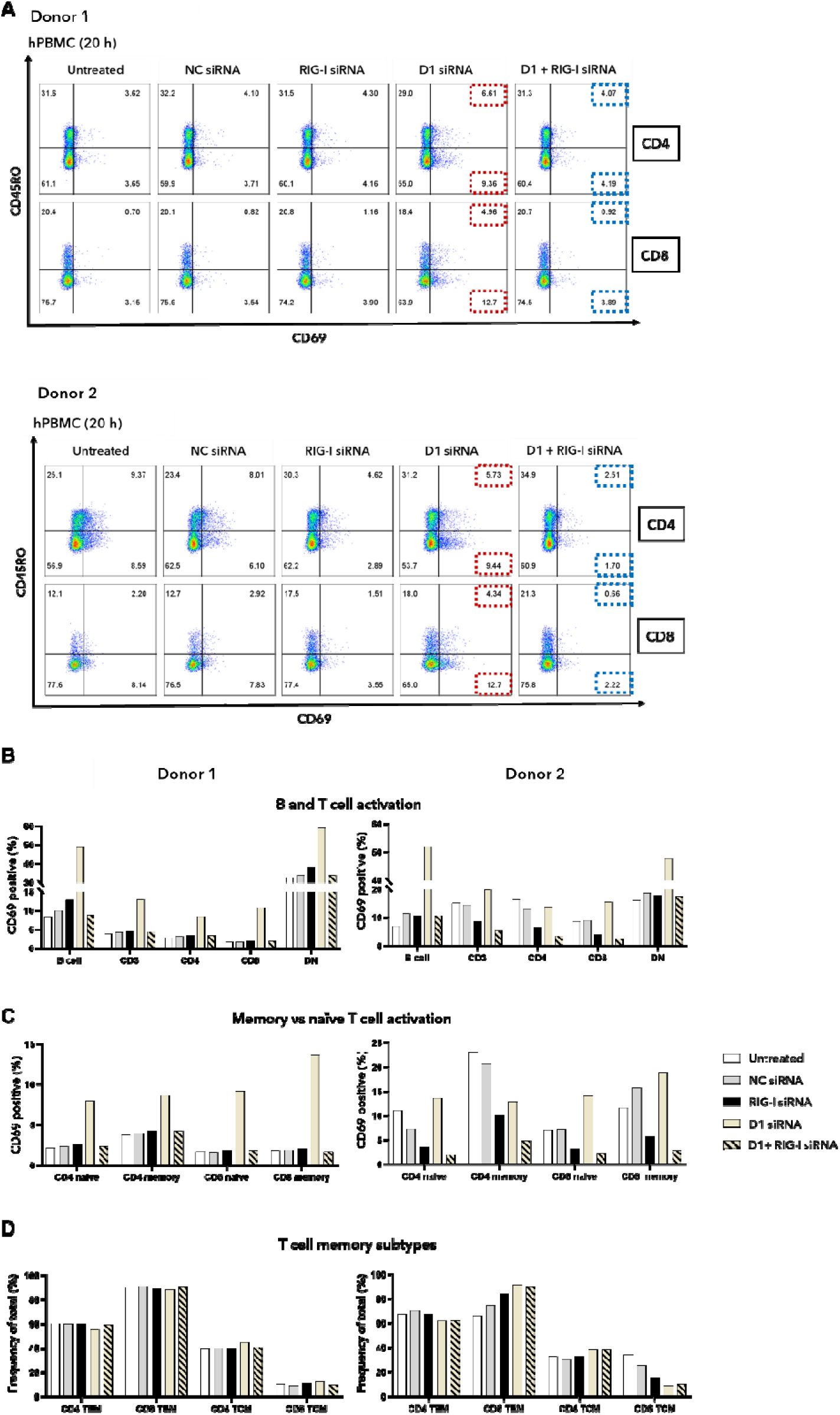
Cytokines produced by lung adenocarcinoma cells upon DUSP11 knockdown activate peripheral blood cells. (A-B) hPBMCs from two separate healthy donors were incubated for 20 h with cell-free media from H1975 cells treated with 10 nM of the reported siRNAs for 48 h, then cell type/lineage and activation status were assessed by antibody staining and analyzed by flow cytometry. (**A**) Pseudocolor dot plots showing activation of CD45RO+ (memory) and CD45RO-(naïve), CD4+ (*top row*), and CD8+ (*bottom row*) T cells as determined by CD69 positivity. (**B**) Activation of various cells types as determined by CD69 positivity (top). B cells (CD3-lymphocytes), CD4+ T cells, (CD3+/CD4+), CD8+ T cells (CD3+/CD8+), and double negative T cells (CD3+/CD4-/CD8-lymphocytes) were assessed. (**C**) Activation of T cell subtypes as determined by CD69 positivity. Naïve (CD45RA+/CD45RO-) and memory (CD45RA-/CD45RO+) CD4+ and CD8+ T cells were assessed. For B-C, the percentage of CD69 positive cells relative to total cell population for each donor is shown on the *y*-axis. (**D**) Percent of memory T cell subtypes (T_CM_: CD45RO+/CCR7+; T_EM_: CD45RO+/CCR7-). The percentage of memory subtypes relative to total CD4 or CD8 memory T cells for each donor are shown on the *y*-axis. For both donors, both B cells and T cell subpopulations exhibited increased activation when incubated with media from D1 siRNA treated cells compared to cells treated with control siRNAs or co-treated with D1 and RIG-I siRNA. Representative gating schemes are shown in Figure S10.

### Mechanisms of DUSP11 modulation may be generalizable to other cancer types

Correlative patient data suggest that other cancer types may be susceptible to DUSP11 modulation (Figure S1). As the DUSP11/3pRNA/RIG-I axis is conserved across nearly all cell types and both proteins are ubiquitously expressed, it is plausible that the immune mediated anti-cancer mechanism elucidated for LUAD could be generalizable to other cancer types or subtypes. Therefore, we evaluated the impact of DUSP11 knockdown on cell viability of various other cell lines to determine what cell types or subtypes could be susceptible to DUSP11 knockdown (i.e., identifying responders versus non-responders). The cell types/subtypes screened included non-cancerous embryonic kidney (HEK293FT; **Figure S2B**), breast cancer (BT474, SKBR3; **Figure S2C**), liver cancer (HepG2, HUH7.5.1; **Figure S2D**), lymphoma (SUDHL4, Ramos; **Figure S2E**), and leukemia (NALM6, MEC1; **Figure S2F**) cells. Compared to controls, a single treatment with 10nM D1 siRNA decreased cell viability of breast cancer cell lines (BT474, SKBR3; **Figure S2C**) when assessed at 96 h; however, no differences were observed in either liver, leukemia, or lymphoma cell line (**Figure S2D-F**). Interestingly, HEK293FT cells did not exhibit a decrease in cell viability when transfected with D1 or D2 siRNA but did show decreases for controls and D3 siRNA (**Figure S2B**). This variable and non-specific decrease in cell viability could indicate that HEK293FT cells are prone to RNA induced toxicity, likely mediated by immune recognition of certain double stranded (ds) RNA sequences, rather than from the knockdown of DUSP11.

The lack of response upon challenge with DUSP11 siRNA may arise from differences in the ability of each cell line to be transfected, resulting in insufficient intracellular delivery of siRNA and subsequent target knockdown. Therefore, we tested this hypothesis in a subset of LUAD, liver, leukemia, and lymphoma cell lines by transfecting 10 nM non-targeting siRNA labeled with a fluorescent dye (TYE™ 563-siRNA) for 24 h and assessing transfection efficiency using the percentage of TYE™ 563 positive cells after flow cytometric analysis. Transfection efficiencies for all cell lines was ∼100% except for MEC1 (∼60%) and Ramos (∼40%); however, the MFI of cells transfected with TYE™ 563-siRNA relative to cells transfected with NC siRNA differed between cell lines (**Figure S3A**). To determine if differences in MFI represented a functional difference in knockdown of our target, we assessed the mRNA expression of *DUSP11* in a subset of these non-responder cancer cell lines (A549, HepG2, Huh7.5.1) 24 h after 10 nM D1i siRNA transfection. Compared to responder cell lines (H1975, H3255), no significant difference in DUSP11 expression was noted (**Figure 1B and S3B**) suggesting that, for the subset tested, MFI did not correlate with the relative expression of DUSP11. Because the leukemia and lymphoma cell lines exhibited the lowest MFI of all cells, and because MEC1 and Ramos exhibited lower overall transfection efficiency, further analysis is still needed to determine whether *DUSP11* mRNA levels were similarly knocked down in these cell lines.

Finally, to determine whether the mechanism of cell death in breast cancer was similar to that in LUAD, we treated SKBR3 cells with 10 nM NC or D1 siRNA in the presence or absence of 10 nM RIG-I siRNA and evaluated the impact on cell viability as before. Like LUAD cells, RIG-I knockdown recovered the cell viability phenotype observed upon DUSP11 knockdown when assessed at 96 h (**Figure S9**). These data support that the immune mediated mechanism via alteration of the 3pRNA pool and sensing of 3pRNA by RIG-I, as observed in LUAD, is likely conserved in breast cancer.

## Discussion

In 2025, lung cancer will continue to account for the majority of cancer-related deaths worldwide(55,56). LUAD, the predominate subtype of NSCLC, contributes significantly to this statistic with 5-year survival rates that have remained under 20% for decades(57). As public health initiatives have significantly decreased tobacco usage rates worldwide, lung cancer prevalence and deaths are expected to drop over the next decades. However, LUAD is not commonly associated with smoking and therefore will remain a significant threat to human lives. Instead, LUAD is often caused by acquired somatic mutations or rearrangements in oncogenes and/or tumor suppressors that lead to dysregulated cellular metabolism and sustained proliferative growth(58,59). Many efficacious targeted therapies have been developed to inhibit these oncogenic proteins, such as those targeting EGFR (Osimertinib, Amivantamab), ALK (Alectinib), ROS1 (Crizotinib), BRAF (Dabrafeib-trametinib), RET (Pralsetinib), and HER2 (Trastuzumab-deruxtecan), among others(60,61). Unfortunately, most patients develop resistance and progressive disease within the first 12-24 months of treatment with these targeted therapies (62), often requiring adjustments to second- or third-line options. The success of ICIs in various other types of cancer brought optimism for their application to LUAD; unfortunately, minimal success has been achieved in oncogene positive LUAD(63), mainly attributed to its ‘cold’ or immunosuppressive TME with low mutational burden (mutational burden is typically associated with smoking, or carcinogen exposure of lung cancer). Further complicating treatments, a significant increased risk of irAEs has been shown when ICIs are administered in combination with or temporally close to TKI administration, especially for EGFR positive cancers(64,65). These two issues – acquired resistance and lack of effective alternative treatments – present a clear unmet need for novel methods to target and treat LUAD and suggest methods to establish a ‘hot’ or inflammatory TME could be efficacious.

Cancers have adopted various ways to mask their immunogenicity and avoid immune destruction, including subversion of immune checkpoint proteins that, under homeostasis, normally prevent the immune system from overreacting and killing healthy cells. The transmembrane proteins, programmed cell death protein 1 (PD-1) and cytotoxic T lymphocyte antigen 4 (CTLA-4), are classic examples of immune checkpoint proteins in that they trigger immunosuppressive signaling and limit immune responses upon binding to their ligands, PD-L1 and CD80/CD86, respectively (66,67). While these proteins play an important role in maintaining homeostasis (e.g., T cell contraction after infection); their aberrant expression or function can cause diseases such as autoimmunity, chronic infection, and cancer(67,68). Many other proteins play a similar role in helping maintain innate and adaptive immune system homeostasis; however, their role has largely been elucidated in the context of viral infection. One such protein is the atypical intracellular phosphatase, DUSP11, which can hydrolyze viral and host 3pRNA, a pathogen associated molecular pattern (PAMP), and in so doing can minimize RIG-I mediated immune detection and destruction. To date DUSP11’s role as an immunomodulating protein in cancer has not been described; however, based on correlative patient data (**Figure 1A and S1**), we hypothesized that the upregulated expression of DUSP11 in some cancers is an effort to evade immune destruction(20), analogous to cancer-associated upregulation of PD-L1.

Here we describe DUSP11 as an innate immune checkpoint (iIC) protein in lung adenocarcinoma, specifically acting as a mediator between RIG-I and its immunogen, 3pRNA. We demonstrate that the expression of DUSP11 is important for the viability of at least two cancer cell lines *in vitro* and for LUAD tumor engraftment and/or growth *in vivo*. In LUAD, we show that this decrease in survivability is due to induction of apoptosis and that this phenotype depends upon the expression of RIG-I, but not other intracellular PRRs. In addition, we show that innate immune response mechanisms such as type I IFN production and upregulation of ISG expression are stimulated upon DUSP11 knockdown in LUAD and that both depend upon the expression of RIG-I. These findings, in addition to other reports of DUSP11’s role in cancer, could have various implications for the field of IO and are briefly discussed below.

### A broader picture of DUSP11 in cancer

For the first time, we describe a direct immunosuppressive and pro-neoplastic role for DUSP11 in cancer, a mechanism distinct from those previously proposed. In addition, while previous reports noted an association between DUSP11 and cancer, ours is the first study to attempt to manipulate DUSP11’s activity utilizing methods amendable to clinical translation (i.e., gene silencing) for cancer therapy. Our new mechanistic model aligns with previous findings and provides a broader understanding of DUSP11’s role in cancer and suggests opportunities for advancing therapeutic modulation of DUSP11. First, 3pRNA, the proposed immunogen in our work, has been studied extensively as an immunogen in the context of viral infections. However, only a small number of cancers are caused by viruses (none of which include LUAD or breast cancer), and an exceedingly smaller subset of these cancer-causing viruses can introduce 3pRNA into the cell during their lifecycle. Instead, the main driver of 3pRNA accumulation in cancer is likely dysregulated cellular energetics, specifically the production of transcripts by an overactive RNAPIII which has been increasingly linked with cancer progression (21–23,69) and its elevated expression also correlates with poorer patient survival for multiple cancer types (TCGA). RNAPIII has been shown to be responsible for transcribing various RIG-I activating oligonucleotides(15,70), including but is not limited to U6 snRNA, Y RNAs, pseudogene of 5S rRNA (RNA5SP141), short interspersed nuclear elements (SINEs; known as Alu elements in humans), and non-coding vault RNAs (vtRNAs)(50). We speculate that one or multiple of these 3pRNAs, such as the vtRNAs recently described in crosstalk with RLRs(51) or upregulated in PDAC(26), are contributing to the activation of innate immune mechanisms described when DUSP11 is knocked down in LUAD. Determining which transcripts are promoting RIG-I activation could have implications in therapeutic development and in improving the efficacy of targeting iIC proteins like DUSP11. Similarly, 3pRNA based reagents have shown minimal success in clinical trials as monotherapy and this is often attributed to toxicity and to complexities brought on by underdeveloped RNA delivery methods in a clinical setting. Modulating DUSP11 expression or activity in an adjuvant setting (**Figure 5**), especially in a tumor-targeted manner, may be advantageous so that efficacious anti-cancer responses can be achieved at a lower administered dose of 3pRNA. This approach may be especially useful for cell types that do not respond to modulating DUSP11 expression alone (**Figure S2**). Second, because DUSP11 can act upon and alter the expression of various 3pRNAs, including many ncRNAs with unknown functions, it is feasible that there are likely multiple cancer type or subtype specific anti-cancer and pro-cancer mechanisms at play aside from the more generalizable innate mediated mechanism we describe here such as the one described for gem resistance in PDAC(26). In addition, given that DUSP11 can alter RNA pools and that AZIN1-AS1 was described to indirectly modulate DUSP11 expression(25), a mechanism involving reciprocal regulation of these two agents could promote cancer survival by keeping immune stimulation at bay, although further studies are needed to confirm or deny this speculation. Finally, DUSP11 expression and activity has been shown to be modulated by various targetable mechanisms, including the activity of the p53 tumor suppressor(24) and the expression of lncRNAs and/or miRNAs(25). Aside from targeting DUSP11 directly, as we have done using siRNA, these targets provide alternative opportunities to further modulate DUSP11 activity as an iIC.

### Implications for cancer immunotherapy

Anti-tumor immunity can be engaged through a variety of mechanisms including the activation of PRRs and subsequent immunostimulatory cytokine release(6,7). Type I IFNs directly regulate the transcription of over one hundred genes that can promote a myriad of cell-intrinsic anti-cancer effects(34,38,71). Type I IFNs have also been shown to have various indirect immunomodulatory effects on both the innate and adaptive immune system, including but not limited to: (i) upregulation of MHC molecules and antigen presentation on a variety of cell types, (ii) recruitment and promotion of effector and memory functions of cytotoxic T cells, (iii) proliferation and increased effector function of NK cells, and (iv) limiting effects of immunosuppressive T regulatory cells and myeloid-derived suppressor cells (34,71). Supporting these claims, we demonstrate that in LUAD cell lines, DUSP11 knockdown greatly increased the release of IFNβ (and potentially other cytokines not evaluated) that potently activated ISRE reporter cells (**Figure 3**) and B and T cell populations from hPBMCs (**Figure 6**) in a paracrine fashion.

In addition to type I IFN release, we also showed that multiple ISGs, including *ISG15* and *IFIT1*, were significantly upregulated upon DUSP11 knockdown. ISGs also regulate the transcription of many genes, for which their protein products can have either anti-tumor or pro-tumor effects on cancer immunogenicity and on the cancer cells themselves, dependent on the cancer type (71–73)(interferome.org). Both IFNβ and secreted ISG15 have been implicated in the transcriptional regulation of MHC proteins in multiple cell types and in promoting NK cell activation and maturation (71,73). Although minimal changes were observed in circulating cell populations in our mouse model, NK cells did show upregulation of MHC class II molecules, and this upregulation was specific to mice that received cells treated with D1 siRNA (compared to NC siRNA). While little is known about the effect of MHC class II expression on NKs, understanding the interactions that caused these changes may be important to define the immunomodulatory effects of modulating DUSP11 expression. However, as mentioned above, T cell deficiency and limited cross reactivity between human cytokines and mouse receptors in this mouse model raise the question of whether these changes are conserved in immunocompetent mice. It will be important in future studies to establish whether tumor infiltrating immune cells are recruited by signaling from tumor cells in which DUSP11 has been knocked down.

Taken together, our data suggests that RIG-I activation and subsequent inflammation are suppressed by DUSP11 activity in LUAD, and that therapeutic intervention to modulate DUSP11’s function could induce broad inflammation in the TME. However, as many of the inflammatory factors mentioned above have also been implemented in disease progression (suggesting there is a balance or tipping point), it remains a priority to better define the type and amount of inflammatory factors released upon DUSP11 modulation. This would help us not only understand their impact on cancer intrinsic properties and immune effector cells (i.e., function), but also calibrate treatments to balance anti-tumor and pro-tumor effects, either alone or in combination with other therapeutics (e.g., TKIs and ICIs). For example, Type I IFN induced immunity has been shown to be important for efficacy of chemotherapy and targeted drugs (74) and for overcoming resistance to immunotherapies(37). In alignment with this, we note that RIG-I knockout decreased sensitivity of H1975 cells to the EGFR TKI, Osimertinib (**Figure 4G)**, which correlates well with another report that IFN responses are induced upon treatment with Osimertinib and are associated with a positive clinical response (75). It is thus possible that RIG-I (and DUSP11 activity) mediates such a response, although this potential mechanistic link requires further evaluation. Overall, it will be crucial to test the efficacy of modulating DUSP11 expression or activity on already growing or metastasized tumors and prioritizes development of clinically useful reagents and delivery methods that enable effective targeting of DUSP11 in specific tissues *in vivo*. For example, one such approach for targeted delivery of siRNA is the use of aptamer technology for which we(76,77) and others(78,79) have shown can effectively localize payloads to cancer cells *in vitro* and to tumors *in vivo*. Alternatively, local delivery methods could be explored such an aerosol inhalation or intranasal drip for delivery of siRNA to the lungs or CNS(80,81).

### Implications for cancer immunoprevention

Aside from cancer immunotherapy, an emerging focus of the IO field is that of cancer immunoprevention, which seeks to modulate the immune system to prevent the development of cancer. In alignment with this goal, the NCI has developed the Cancer Immunoprevention Network (CIP-Net, https://www.prevention.cancer.gov) that supports research focused on the discovery of novel immunoprevention targets and pathways. While much of this work has focused on utilizing the adaptive immune system(82,83), increasing evidence indicates that there may be an adjuvant or potentially separate role for pathways and cell types associated with innate immunity. Here we showed that pre-treating LUAD cells with a single 10 nM dose of DUSP11 siRNA prevents or slows tumor engraftment and/or subsequent tumor growth. This finding, along with other supporting data shown here, suggests that modulating the expression and activity of iIC proteins such as DUSP11, (like traditional ICIs) prior to development of cancer, could potentially create a cell intrinsic and extrinsic environment that is less compatible with tumor cell growth, development, or metastasis. In principle, a localized application could provide advantages, as DUSP11 is ubiquitously expressed at variable levels in all tissues, and its decreased activity could potentially increase susceptibility to autoimmunity. However, DUSP11 knockout mice are viable (like PD1 KO mice), and our data suggest that toxicity of DUSP11 knockdown may be limited in non-cancerous cells, suggesting that systemic inhibition could be feasible in some cases. Nevertheless, future studies should include evaluations of whether DUSP11 is indeed a good target for cancer immunoprevention, whether alone or as an adjuvant to reagents currently in clinical trials.

### Predicting cancer types that might respond to modulating DUSP11 activity

Published reports and patient data show that DUSP11 expression correlates with survival for many cancer types. Indeed, we showed that decreased expression of DUSP11 decreases LUAD cell viability and tumor growth. However, not all correlations were necessarily predictive as we have shown for breast cancer (SKBR3, BT474), liver cancer (HepG2, Huh7.5.1), and one LUAD (A549) cell line (**Figure S2**). Two limitations in our work – cell susceptibility screening model and bioinformatic analysis methods – provide guidance to help better elucidate the cell populations for which DUSP11 modulation could be an effective treatment. First, because there are both cell-autonomous effects and paracrine mediated effects of DUSP11 knockdown, future cancer cell type susceptibility screens should utilize a model that also harbors an intact immune system (e.g., organ on chip or mouse model) as opposed to the MTS assay used here which only evaluates cell-autonomous effects. Second, as cancer is readily defined as a heterogenous disease, when possible during bioinformatic analysis, (i) the sample size and type should be increased (e.g., including samples from more diverse patient populations), (ii) the data should be further differentiated by subtype and stage (similar to what was done for iCCH(19)), and (iii) the datasets should be validated. In regards to the latter, the Human Protein Atlas recently uploaded validated breast cancer sample data (accessed January 2025) which now shows an opposite correlation from TCGA data and is more in line with our findings in **Figure S2C**. In addition and because there are many variables at play (i.e., DUSP11, RNAPIII, 3pRNA, and RIG-I), we suggest that a more in depth evaluation of a biomarker ‘panel’, which assesses the expression and/or activity of all variables, may provide a more biologically relevant scenario and could improve predictive value.

## Conclusion

Our preclinical studies suggest that DUSP11 is an excellent target to modulate innate immunity and induce cell death in LUAD. As DUSP11 acts to suppress the immune response (checkpoint), removing this ‘brake’ generates an anti-cancer and immunostimulatory environment (**Figure 7**). While early data suggest that this target and mechanism may be generalizable to other cancer types, including breast, it will be important to understand better the mechanisms that lead to apoptosis upon the modulation of DUSP11 expression, to identify which cancer types respond to DUSP11 inhibition, and to identify associated biomarkers. Considering our findings, the development of clinically feasible methods to target this protein and assessment of the safety and toxicity of such targeting warrants further investigation.

**Figure 7.**
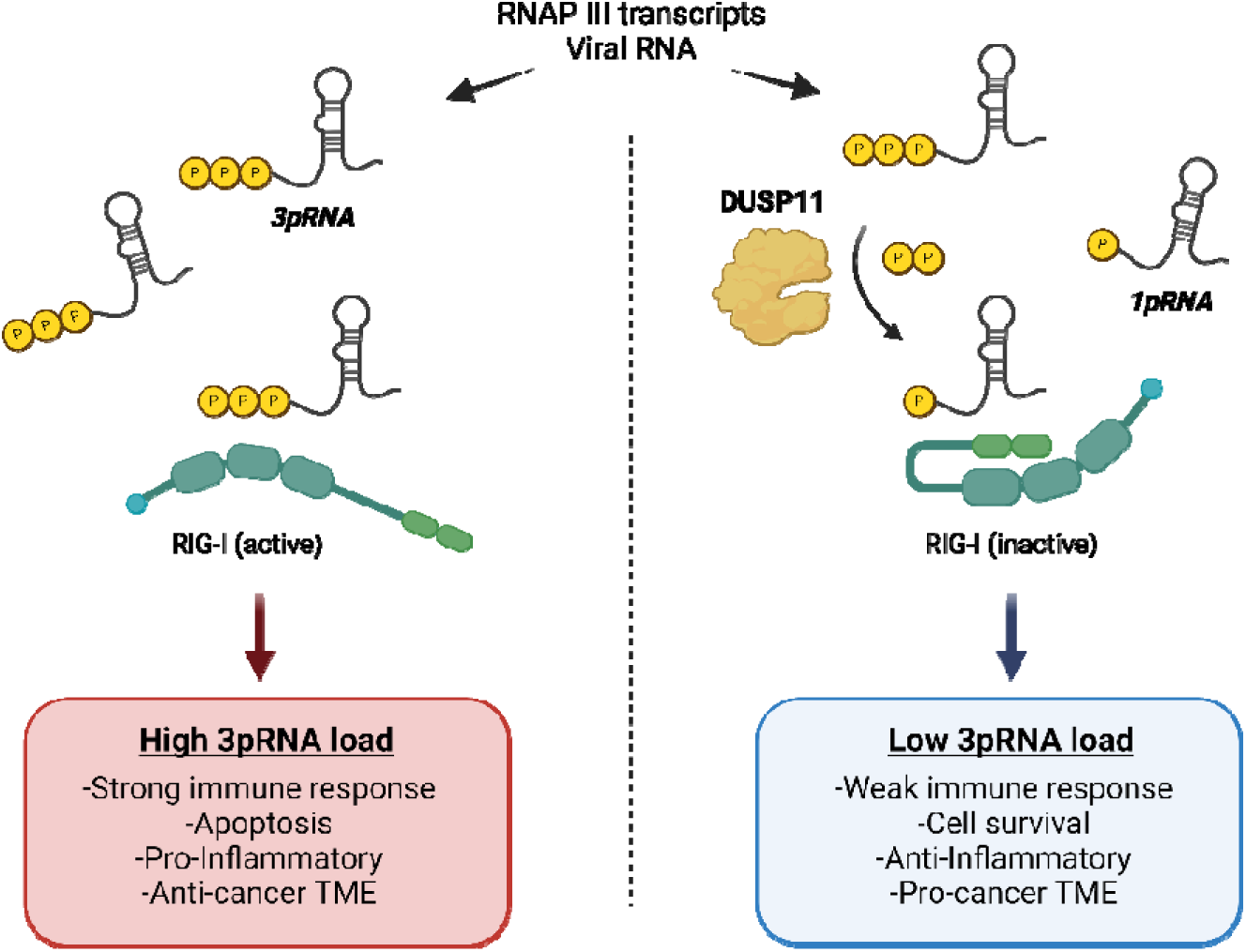
Model of DUSP11’s immunosuppressive role in cancer. Schematic showing DUSP11’s immunosuppressive role in cancer. 5’ triphosphate RNA (3pRNA) transcripts are produced predominately by RNA polymerase III (RNAP III) in cancer or by viral polymerases during infection. In the presence of DUSP11, 3pRNA is converted to 1pRNA, which is not recognized by RIG-I, an intracellular PRR, thus avoiding triggering an immune response and creating a pro-neoplastic environment. In the absence of DUSP11, 3pRNA is not converted and is recognized by RIG-I, which in turn triggers an innate immune response that contributes to an inflammatory and anti-cancer environment. Graphic made using Biorender.com

## Materials and Methods

### Reagents

DNA and RNA oligonucleotides were purchased from Integrated DNA Technologies (IDT, Coralville, IA). Primary antibodies for probing western blots were purchased from Cell Signaling Technology (Danvers, MA; RIG-I, cat #3743, RRID:AB_2269233), Proteintech (Rosemont, IL; DUSP11, cat #10204-2-AP, RRID:AB_2277484), BioLegend (San Diego, CA; GAPDH, cat #607902, RRID:AB_2734503), and Santa Cruz Biotechnology (Dallas, TX; β-actin, cat #sc-47778 HRP, RRID:AB_2714189). Secondary antibodies for western blots were purchased from Invitrogen (Waltham, MA): anti-Mouse (cat #G-21040, RRID:AB_2536527), anti-Rabbit (cat #G-21234, RRID:AB_2536530), anti-Rat (cat #A18871, RRID:AB_2535648). Dual color protein ladder standards were purchased from Biorad (cat #1610374). BSA standards and Pierce detection reagents were purchased from ThermoFisher Scientific (Waltham, MA). Protease and phosphatase inhibitor cocktails and RIPA buffer were purchased from Abcam (Cambridge, MA). All other reagents and materials, including chemotherapeutics and small molecule inhibitors, were purchased from Sigma Aldrich (St. Louis, MO) unless otherwise noted.

### Cell culture

A549 cells were gifted by Dr. Bumsuk Hahm, University of Missouri-Columbia. H3255, H1975, and H820 were gifted by Dr. Raghuraman Kannan, University of Missouri-Columbia. SKBR3 and BT474 cells were gifted by Dr. Thomas Quinn, University of Missouri-Columbia. All other cell lines were purchased by ATCC (Manassas, VA). A549 (RRID:CVCL_0023), SKBR3 (RRID:CVCL_0033), BT474 (RRID:CVCL_0179), HEK293FT (RRID:CVCL_6911), and HepG2 (RRID:CVCL_0027) cells were cultured in DMEM supplemented with 10% Fetal Bovine Serum (FBS; Gibco, ThermoFisher Scientific), 1 mM sodium pyruvate, 2 mM L-glutamine, and 1X NEAAs. H1975 (RRID:CVCL_1511), H3255 (RRID:CVCL_6831), H820 (RRID:CVCL_1592), Huh7.5.1 (RRID:CVCL_E049), NALM6 (RRID:CVCL_0092), MEC1 (RRID:CVCL_1870), SUDHL4 (CVCL_0539), and Ramos (RRID:CVCL_0597) cells were cultured in HyClone™ RPMI 1640 supplemented with 10% FBS, 1 mM sodium pyruvate, 2 mM L-glutamine, and 1X NEAAs. All cell lines were maintained at 37°C in humidified incubator with 5% CO_2_ and passaged when ∼80% confluent. Cells were not passaged more than 20 times from thaw or past total passage number 30 (P30) to minimize genetic drift. Consistency in cell morphology was checked prior to each passage. For lung cancer cells expressing EGFR, sensitivity to Osimertinib (TKI) was checked regularly and compared to previous observations.

### Cell transfection

Three siRNA transfection methods were tested using a fluorescent dye labeled siRNA. Relative transfection efficiency and MFI in HEK293FT, H3255, and H1975 cells, as measured by flow cytometry, was Lipofectamine^TM^ RNAiMAX > Fugene® SI > > jetPRIME® = polyethylamine (PEI). Therefore, for subsequent assays requiring siRNA treatment, cells were transfected using Lipofectamine^TM^ RNAiMAX (Invitrogen, cat #13778150,) per manufacturer protocol. In brief, siRNA and Lipofectamine reagent were separately diluted to 2X desired concentration in RPMI supplemented as above (*see M&M section on cell culture*) but with 0% FBS. Equal volumes of the two were mixed and then allowed to assemble into functional complexes for ∼10min before adding to cells. For co-transfection experiments, the siRNAs or 3pRNA (InvivoGen, San Diego, CA; cat #tlrl-hprna) were mixed and diluted to 2X desired concentration in RPMI before being mixed and complexed for ∼10min with the transfection reagent. For co-transfection experiments, all transfected RNA concentrations were kept consistent using filler RNA (NC siRNA) to minimize misinterpretations. All Dicer substrate siRNA sequences are shown in Table S1.

### Cell viability assays

Cells were seeded at ∼5000 or 10,000 cells/well in a 96-well flat-bottom tissue culture plate 24 h prior to treatment. Cells were treated (transfected) with siRNA at reported concentrations or left untreated for 48 h (48 h timepoint) and media was replaced for an additional 48 h (96 h timepoint) before adding MTS-based cell viability reagent (CellTiter96, Promega, Madison, WI). MTS-reagent was allowed to incubate for ∼2 h at 37°C before obtaining absorbance values at 490 nm. Cell viability is reported relative to untreated samples. For dose response experiments, EC50 values were determined by fitting a nonlinear curve for inhibition using a least squares regression method (no weighting) using GraphPad Prism v10.4.0 for Windows (GraphPad Software, Boston, MA).

### Cytotoxicity assay

Cells were seeded at ∼15,000 cells/well in a 48-well flat-bottom tissue culture plate 24 h prior to treatment. Cells were treated with siRNAs at reported concentrations for 48 h and stained (48 h timepoint) or media was replaced for an additional 48 h prior to staining (96 h timepoint). Briefly, cells were moved to a 1.5 mL Eppendorf tube, resuspended in Live Cell Imaging Solution (Invitrogen), and stained with CellEvent™ Caspase-3/7 Green Detection Reagent (ThermoFisher Scientific, cat: #C10431) for 25 min and Helix NP™ NIR (BioLegend cat: # 425301) Dead Cell Stain for 5 min at 37°C and 5% CO_2_. Cells were immediately analyzed via flow cytometry.

### Western blots

Cells were seeded at ∼100,000 cells/cm^2^ in a 12-well flat-bottom tissue culture plate 24 h prior to treatment. Cells were treated with siRNA at the reported dose for 24 h. Cells were washed with ice cold 1X DPBS and lysed in 1X RIPA buffer supplemented with phosphatase and protease inhibitor cocktails on ice for 30 min (with scraping). Cell lysates were either kept on ice and used immediately or stored at -80°C. Prior to gel loading, lysates were quantified using Pierce^TM^ 660nm Protein Assay Reagent (ThermoFisher Scientific cat #22660) and then ∼4 µg of total cell lysate in 1X Laemmli loading buffer were loaded into a 4-12% SurePAGE gel (GeneScript, Piscataway, NJ). PAGE gels were run at 130V for ∼1h in Bis-Tris buffer (GeneScript) and then wet transferred to a 0.2 µm PVDF membrane (BioRad, activated in 100% methanol) at 100V for ∼1 h in 1X transfer buffer containing 20% methanol. Blots were blocked for 1 h at RT in 1X Tris-buffered saline with 0.1% Tween 20 (TBST) + 3% BSA Fraction V (Gold Biotechnology, Olivette, MO) and then incubated in primary antibody diluted in blocking buffer with overnight rocking at 4°C (total DUSP11 and RIG-I) or for 1 h at RT (GAPDH and β-actin). Blots were washed for 5-10 min in 1X TBST (x3) and incubated in secondary antibodies diluted in blocking buffer for 1 h at RT. Blots were washed again in 1X TBST (x3) and then incubated with SuperSignal^TM^ West Pico PLUS Chemiluminescent substrate (ThermoFisher Scientific) in the dark for 5 min prior to imaging on an Invitrogen^TM^ iBright^TM^ 1500 imaging system. Blots were stripped using two 5-10 min washes with mild stripping buffer (pH = 2) followed by two 15 min washes with 1X TBST. After stripping, blots were blocked and re-probed as above. All images were analyzed using FIJI (ImageJ; RRID:SCR_003070). All quantified bands for phosphorylated and total protein were normalized to GAPDH signal prior to plotting.

### RT-qPCR and analysis

Cancer cells were seeded at ∼100,000 cells per well in 12-well plates and incubated overnight before treatment with reported concentration of siRNAs for 24 or 48 h. Then total RNA of cells was isolated by Total RNA miniprep kit (New England BioLabs, cat #T2010S) per manufacturer protocol. RNA was reversed transcribed using Takara SMART MMLV RT (cat #S4989, Takara) and amplified using iTaq Universal SYBR Green Supermix (cat #1725121, Bio-Rad), and imaged on a BioRad qPCR system. Analysis was done using BioRad CFX Matero (v4.0.2325). All RT-qPCR primer sets are shown in Table S2.

### Peripheral blood cell isolation, assay, and staining

All fluorescent dye labeled antibodies were purchased from BioLegend unless otherwise noted. Human peripheral blood mononuclear cells (hPBMCs) were stained with the following antibodies: CD3 (AF700, cat #300424, RRID:AB_493741), CD4 (SB550, cat #344656, RRID:AB_2819979), CD8 (APC/Fire810, cat #344764, RRID:AB_2860890), CD45RO (PE, Invitrogen, cat #12-0457-42, RRID:AB_1272079), CD45RA (PE-Cy7, cat #983006, RRID:AB_2922655), CD69 (APC, cat #310910, RRID:AB_314845), and CCR7 (PE-Cy5.5, Invitrogen, cat #35-1979-42, RRID:AB_2815128). Mouse peripheral blood cells were stained with the following antibodies: CD11c (PE, cat #117308, RRID:AB_313777), F4/80 (APC/Cy7, cat #123118, RRID:AB_893477), NK1.1 (PE/Cy7, cat #108714, RRID:AB_389364), MHC II I-A/I-E (AF700, cat #107622, RRID:AB_493727), CD69 (APC, cat #104514, RRID:AB_492843), CD86 (FITC, cat #105006, RRID:AB_313149), Ly6c (PerCP/Cy5.5, cat #128012, RRID:AB_1659241).

#### For mouse peripheral blood

At 24 h and 7 days post tumor cell implantation, mice tail veins were bled and ∼30 µl of blood was extracted and immediately mixed with heparin to prevent coagulation. Red blood cells (RBCs) were then lysed on ice for ∼5 min with ACK (ammonium-chloride-potassium) buffer and centrifuged at 500xg for 5min to remove lysed RBCs. Cells were resuspended for 20 min in ice cold Fc blocking antibody (BioLegend, cat #156604) diluted in FACS buffer (1X DPBS + 0.5% BSA) for 20 min. Cells were washed with FACS buffer and then resuspended in ice cold antibody solution diluted in FACS buffer for 1 h. Cells were again washed with FACS buffer and fixed in 4% PFA for 10 min at 4°C. Stained and fixed cells were resuspended in FACS buffer and immediately analyzed by flow cytometry.

#### For human peripheral blood

Heparin-anticoagulated peripheral blood was obtained from healthy adults who provided written informed consent in accordance with protocol 2031144 approved by the Institutional Review Board of the University of Missouri. hPBMCs were isolated by Ficoll-Hypaque (Cytiva, Marlborough, MA) density centrifugation and then stored at -80°C in 90% FBS plus 10% DMSO. On the day of the experiments, hPBMCs were thawed quickly and added to warm RPMI 10 at 37°C. Cells were centrifuged at 500xg for 5 min and then resuspended in fresh warm RPMI 10, counted, and ∼100,000 cells were added to an untreated 96-well round bottom cell culture plate. H1975 cells were treated for 48 h with either 10 nM siRNAs or left untreated and centrifuged gently to avoid cell transfer, then cell-free media taken from these cells was added to the hPBMCs at the reported dilutions and incubated for ∼20 h at 37°C. Cells were centrifuged at 500xg for 5 min and resuspended in ice cold Fc blocking antibody (BD Biosciences, San Diego, CA, cat #564219) diluted in FACS buffer for 20 min. Cell were washed with FACS buffer and then resuspended in ice cold antibody solution diluted in FACS buffer for 30 min. Cells were again washed with FACS buffer and fixed in 4% PFA for 10 min at 4°C. Stained and fixed cells were resuspended in FACS buffer and immediately analyzed by flow cytometry.

### Flow cytometry and analysis

Flow cytometry was performed on an Attune NxT (ThermoFisher Scientific, Waltham, MA) by counting a minimum of 10,000-30,000 or 100,000 (peripheral blood) live cell events. For live/dead staining, far-red dyes (HELIX NP™ NIR) were excited using a 637 nm laser, and fluorescence was detected using a standard RL1 filter [670 ± 7 nm]. For CellEvent^TM^ Caspase 3/7 Green Detection Reagent, cells were excited using a 488 nm laser and detected using standard BL1 filter [530 ± 15 nm]. For TYE563 labeled siRNA, cells were excited using a 561 nm laser and detected using standard YL1 filter [585 ± 8 nm]. For peripheral blood staining, fluorescent dye labeled antibodies were either (i) excited using a 488 nm laser and detected using a standard BL1 (FITC, SparkBlue550) or BL3 [695 ± 20 nm](PerCP-Cy5.5, PE-Cy5.5) filter, (ii) excited using a 561 nm laser and detected using standard YL1 (PE) or YL4 [780 ± 30 nm](PE-Cy7) filter, (iii) excited using a 637 nm laser and detected using a standard RL1 (APC), RL2 [720 ± 15 nm](AF700), or an RL3 [780 ± 30 nm](APC-Cy7, APC/Fire810) filter.

Compensation settings were identified from single-antibody staining of cells prior to staining with antibody mixes. Flow cytometry data were analyzed and processed using FlowJo Software v10 (Treestar, Ashland, OR; RRID:SCR_008520). For cytotoxicity studies, singlet cells were gated and analyzed for percent positivity (Caspase 3/7 and Sytox staining). Representative gating strategies for mouse peripheral blood and hPBMC staining are shown in Figure S10.

### RIG-I Knockout Cell Line Engineering

The RIG-I gene was edited through a CRISPR/Cas9 system delivered via lentiviral vector. Oligos for targeting RIG-I (FWD: 5’ caccgTTGGATGCCCTAGACCATGC 3’; REV: 5’ aaacGCATGGTCTAGGGCATCCAAC 3’) were cloned into LentiCRISPRv2 (Addgene#52961, a kind gift from Dr. Feng Zhang) using the Zhang lab protocol(84,85). For lentiviral particle generation, HEK293FT cells were maintained in DMEM supplemented with 2mM L-Glutamine (Cytiva Hyclone) and 7.5% FBS (Gibco). Cells were seeded at ∼500,000 cells per well in a 6-well plate 24 h prior to co-transfected with 450 ng LentiCRISPRv2-RIG-I (a kind gift from Dr. Marc Johnson), 450 ng psPAX2 (Addgene#12260, a kind gift from Dr. Didier Trono; RRID:Addgene_12260), and 100 ng pMD-G (Invitrogen) complexed with 4 μg polyethyleneimine (PEI MW ∼25,000; PolyScience, Inc). After 72 h, virus-containing supernatant was collected and pooled, centrifuged at 1000 x g to remove cellular debris, and transferred to a new tube to be stored overnight at -80°C. The supernatant was thawed on ice, and 8 ml supernatant was mixed thoroughly with 2 ml 50% w/v PEG 8000 (ThermoFisher Scientific), 850 μL 5M NaCl, and 900 μL 1X PBS, and incubated overnight at 4°C. The next day, this mixture was centrifuged at 1500 x g for 45 min at 4°C and the pellet was resuspended in 1 ml PBS. H1975 cells were transduced with viral particles in the presence of 6 μg/ml polybrene (Sigma-Aldrich). After 48 h, cells were transferred to a new plate and selected with 2 μg/ml puromycin (ThermoFisher Scientific) for 1 week, with the media and puromycin being refreshed every 2 days. Single cells from the surviving transduced cells were sorted into individual wells of a 96-well plate by limiting dilution. Clones were allowed to proliferate in the presence of puromycin, and conditioned media was collected from a sub-confluent H1975 parental plate and passed through a 0.45 μm filter. Once proliferated, genomic DNA was harvested, amplified using RIG-I specific primers (FWD: 5’ CGGCCTGGGCAGAGGTTTT 3’; REV: 5’ GCTTCCATTGGGCAGAAATGC 3’), gel extracted, and subject to Sanger Sequencing (University of Missouri Genomics Technology Core) and TIDE analysis(86) to confirm editing. Cell lysates were then subject to western blot to confirm loss of protein.

### *In vivo* tumor burden

To develop cell-line-derived subcutaneous tumor models(87), H1975 cells were grown as described above in 15 cm dishes to ∼50% confluence and then transfected with 10 nM siRNA for 24 h. Cells were washed twice with warm DPBS, detached with TrypLE Express, assessed for viability (see “Cytoxicity” section above) and resuspended in ice cold PBS plus 50% extracellular matrix (ECM) gel (Sigma Aldrich, cat: #E6909) at ∼2.5x10^6^ live cells per 100 µL. Cells were kept on ice during injections to prevent solidification of ECM gel. BALB/c nude mice (Charles River, Wilmington, MA; RRID:IMSR_CRL:194; ∼8 weeks of age) were then briefly anesthetized with isoflurane in an induction chamber (4%, 250 mL/s). Using a 28G 1/2” 1 cc insulin syringe/needle, 2.5x10^6^ cells (100 µL) were subcutaneously injected into the right flank of each mouse. Approximately equal ratios of male to female mice were used for each treatment group. Right flanks were palpated every 2-4 days to check for engraftment, at which point tumors were measured with digital calipers (minimum size ∼4 mm^3^) until at least one mouse reached the humane endpoint (tumor diameter >20 mm or volume >2000 mm^3^). No mice experienced significant weight changes (data not shown). Tumor diameters for width (W) and length (L) were measured, and raw tumor volumes (V; mm^3^) were calculated using the equation: V = 0.5 x L x W^2^. At the end point (day 30 post injection) mice were humanely euthanized and imaged in prone and right recumbent positions. Tumors were excised, *ex vivo* endpoint tumors were imaged, and weights were recorded. No mice were excluded from the analysis. Tumors were placed in 1.5 mL Eppendorf tubes, snap frozen in liquid nitrogen, and stored at -80°C for future analyses.

### Statistical Analyses

All statistical analyses were performed using GraphPad Prism v10.4.0 for Windows (GraphPad Software; RRID:SCR_002798). All statistical analyses are reported in the respective figure legends. For the *in vivo* tumor burden time course, statistical analysis was performed using two methods: (i) a linear regression model and (ii) an area under the curve (AUC) model using a Brown-Forsythe and Welch analysis of variance (ANOVA) with post hoc multiple comparisons test corrected using the Dunnett T3 method. Unless otherwise noted in the figure legend, for all remaining experiments statistical analysis was performed using a one-way ANOVA with post hoc multiple comparisons test corrected using either Šidák (when comparing treatments to multiple controls) or Dunnet’s (when comparing treatment to a single control) methods. For all: *p < 0.05, **p < 0.01, ***p < 0.001, ****p < 0.0001).

## Data availability statement

The data generated in this study are available within the article and its supplementary data files. Raw data generated in this study are available upon request from the corresponding author.

## Animal ethics statement

All animal procedures were conducted according to NIH guidelines for the care and use of laboratory animals and were approved by the University of Missouri Institutional Animal Care and Use Committee.

## Acknowledgements

This work was financially supported by the National Cancer Institute of the National Institutes of Health - Award #F30CA275349 (PI: Thomas) and UM Research and Creative Works Strategic Investment Program Grant (PI: Burke). The authors would like to thank Dr. Haval Shirwan and Dr. Patricia Riberio Pereira for their pre-submission comments.

## CRediT authorship contribution statement

**Brian J. Thomas:** Conceptualization, Methodology, Investigation, Visualization, Formal Analysis, Data Writing – Original Draft, Writing – Review & Editing. **Xue Bai:** Methodology, Investigation, Writing – Review & Editing. **Benjamin J. Cryer:** Resources, Investigation, Writing – Review & Editing. **Sydney M. Escobar:** Resources. **Lee-Ann H. Allen:** Resources, Supervision, Writing – Review & Editing. **Mark A. Daniels:** Resources, Supervision. **Margaret J. Lange**: Resources, Supervision, Writing – Review & Editing. **Donald H. Burke:** Project administration, Resources, Supervision, Writing – Review & Editing, Funding acquisition.

## Synopsis

The protein described in this work (DUSP11) provides a potential new immunotherapeutic target to stimulate the anti-cancer innate immune response and/or enhance anti-cancer adaptive responses. This work also highlights that there are targetable checkpoints within the innate immune system, in addition to those already described for the adaptive immune system.

---

**Figure S1.**
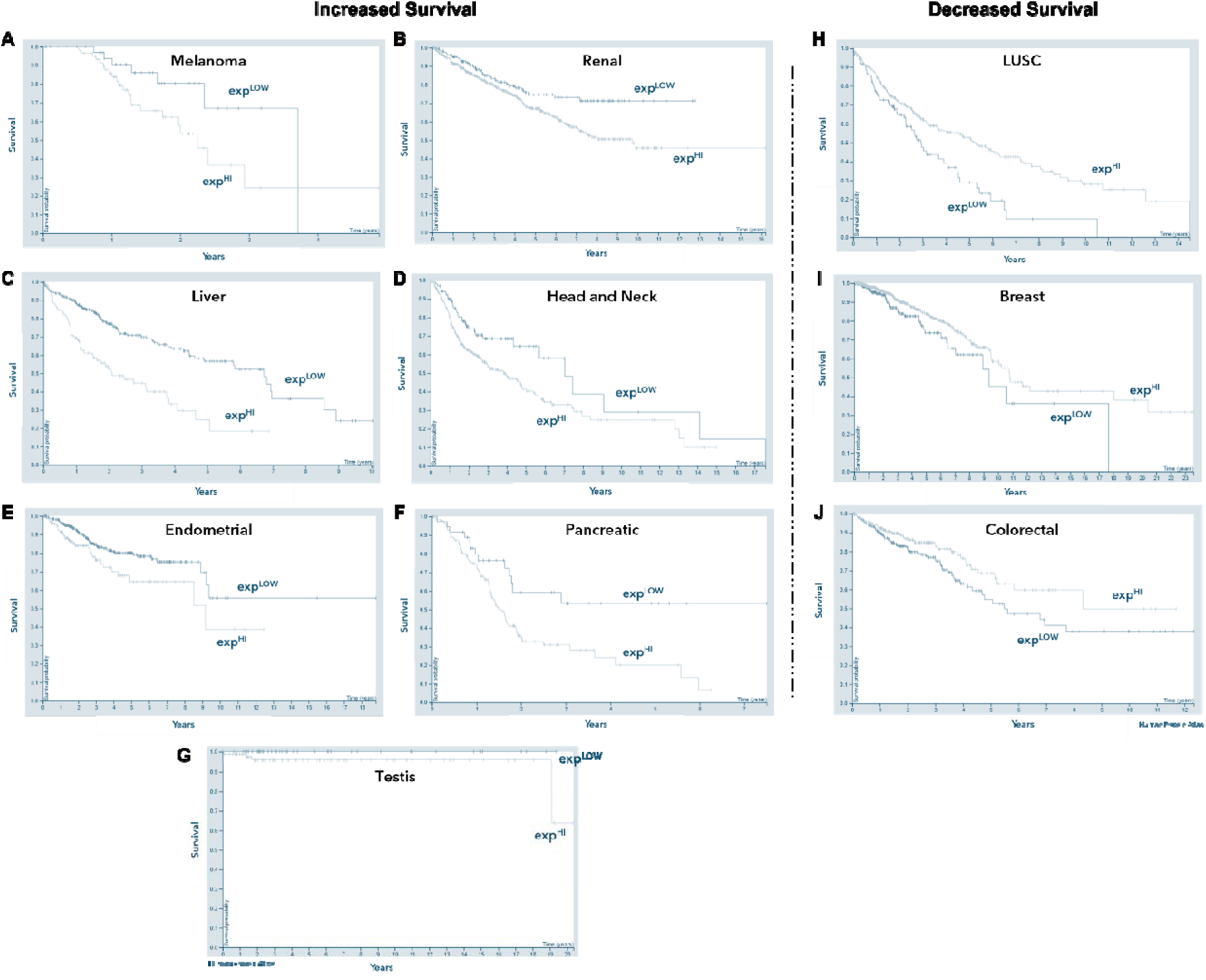
DUSP11 expression is correlated with patient survival for multiple cancer types and subtypes. (A-J) Kaplan-Meier survival curves for patients with multiple cancer types having either improved (A-G) or decreased (H-J) survival correlated with low *DUSP11* mRNA expression. Time (years) is represented on the *x*-axis and percent survival of the population is represented on the *y*-axis. Light blue lines refer to high DUSP11 expression (above TPM cutoff) and dark blue lines refer to low DUSP11 expression (below TPM cutoff). TPM cutoffs used for each cancer type are as follows: (A) melanoma = 12.2, (B) renal = 9.6, (C) liver = 9.2, (D) head and neck = 20.6, (E) endometrial = 7.9, (F) pancreatic = 13.8, (G) testis = 20.3, (H) lung squamous cell carcinoma (LUSC) = 22.5, (I) breast = 16, (J) colorectal = 18.3. Data was accessed from TCGA obtained from the Human Protein Atlas.

**Figure S2.**
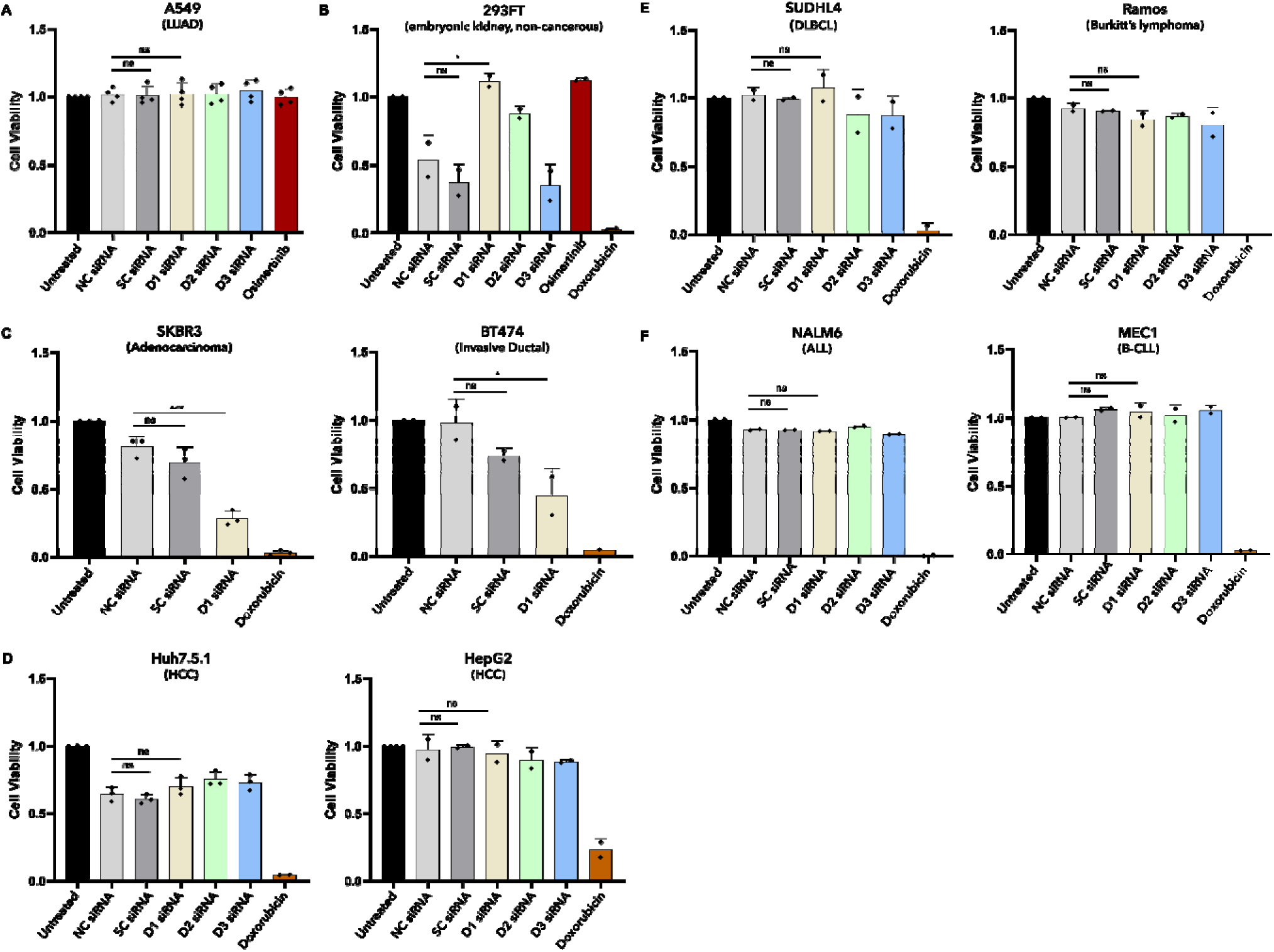
DUSP11 knockdown decreases cell viability of breast but not lymphoma, leukemia, or liver cancer cell lines. (**A-F**) Cell viability was assessed for various cell lines by MTS assay following no treatment (black bars) or 96 h treatment with 10 nM NC (light grey bars), SC (dark grey bars), D1 (sand bars), D2 (green bars), D3 (blue bars) siRNA, 0.5 µM Osimertinib (red bar), or 5 µM Doxorubicin (media change with no siRNA or drug at 48 h). Cell viability relative to untreated cells is reported on the *y*-axis and plotted values represent mean ± SD for n=2-4 independent experiments, each with 2-4 technical replicates. A549 (LUAD, **A**), HepG2, Huh7.5.1 (liver cancer, **D**), SUDHL4, Ramos (lymphoma, **E**), NALM6, and MEC1 (leukemia, **F**) cells exhibited no change in cell viability when treated with D1, D2, or D3 siRNA compared to control siRNAs. **B**) HEK293FT (non-cancerous embryonic kidney cells) exhibited decreased cell viability when treated with NC, SC, and D3 siRNA but not D1 or D2 siRNA, potentially suggestive of an off-targeted immune response to dsRNA rather than to knockdown of DUSP11. **C**) SKBR3 and BT474 (breast cancer exhibited a decrease in cell viability when treated with D1 siRNA but not controls. For all, statistical analyses were performed using a one-way ANOVA with post hoc multiple comparisons test corrected using the Dunnett method (*p < 0.05, ***p < 0.001).

**Figure S3.**
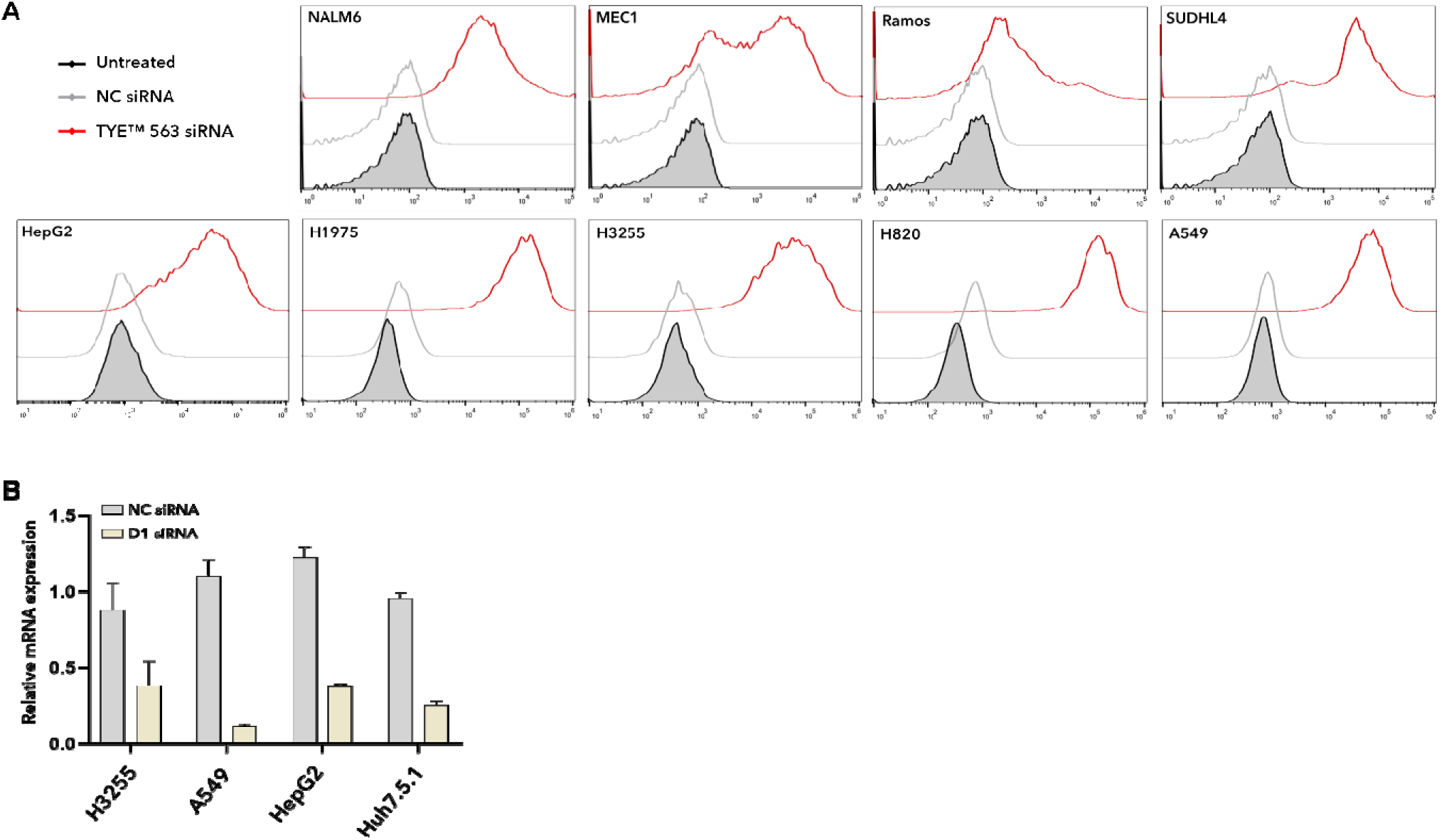
siRNA transfection efficiency and DUSP11 mRNA expression of various cancer cell lines. (**A**) Reported cells lines were left untreated (black line) or transfected with 10 nM NC (grey line) or TYE™ 563 labeled non-targeting siRNA (red line) for 24 h, then transfection efficiency and fluorescence intensity were assessed by flow cytometry. Representative stacked histogram plots with fluorescence intensity (YL1A channel) reported on the *x*-axis from n=1-2 independent experiments. (**B**) Reported cell lines were left untreated or transfected with 10 nM NC (grey bar) or D1 (sand bar) siRNA, and *DUSP11* mRNA was assessed by RT-qPCR at 24 h. All values are normalized to GAPDH and reported relative to mRNA from the untreated sample on the y-axis. Plotted values represent mean ± SD for n=1-2 independent experiments, each with 2-3 technical replicates. These data indicate that non-responder cells lines like A549 and HepG2 were sufficiently transfected and that their target gene was knocked down to similar levels of responder cell lines (H3255, H1975; see Figure 1B).

**Figure S4.**
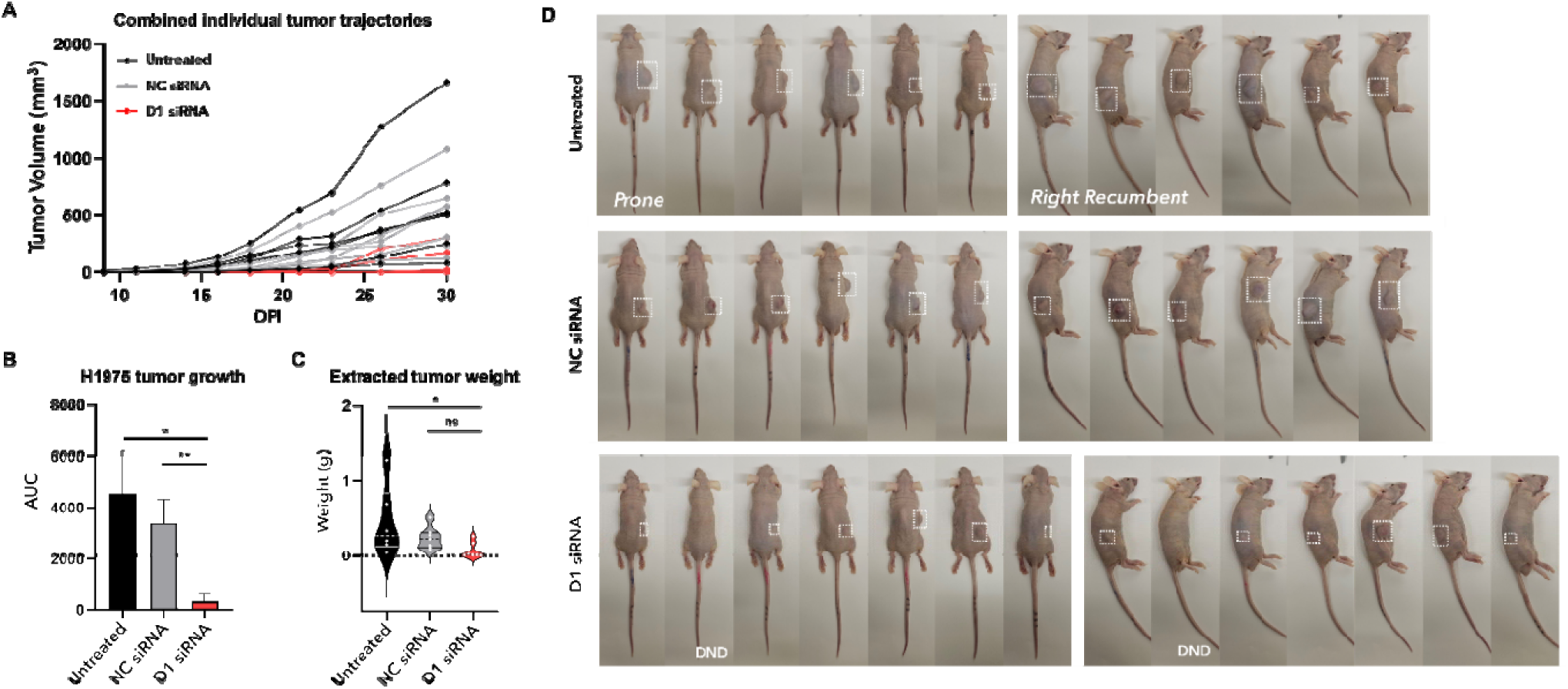
Individual tumor trajectories and endpoint tumor evaluation. (**A-D**) Mice were treated as in the schematic in Figure 2A. (**A**) Combined individual trajectories for n=6 (untreated) or n=7 (NC and D1 siRNA treated) mice from each treatment group (*see* Figure 2C *for trajectories by treatment group*). Days post implantation (DPI) is reported on the *x*-axis and tumor volume (mm^3^ = 0.5 x L x W^2^) is reported on the y-axis. (**B**) Area under the curve (AUC) of tumor growth kinetics, in which the plotted values represent mean ± SD. Using the AUC model, tumor burden from mice implanted with cells treated with D1i siRNA was significantly decreased compared to those implanted with untreated cells or NC siRNA treated cells. (**C**) Violin plots of endpoint *ex vivo* tumor weights of mice from the three treatment groups. Tumor weight is reported on the *y*-axis where the dotted line represents the median and solid line represents the quartiles. Tumors from mice implanted with cells treated with D1i siRNA weighed significantly less when compared to those implanted with untreated cells but not NC siRNA treated cells. (**D**) Mice from each treatment group were imaged in the prone (*left*) and right recumbent (*right*) anatomical positions at the endpoint (DND = tumor did not develop). Note that images for one mouse treated with NC siRNA were not obtained and are not included. For B, statistical analysis was performed using a Brown-Forsythe and Welch analysis of variance (ANOVA) with post hoc multiple comparisons test corrected using the Dunnett T3 method. For C, statistical analysis was performed using a Kruskal-Wallis test with post hoc multiple comparisons test corrected using the Dunn’s method (*p < 0.05, **p < 0.01).

**Figure S5.**
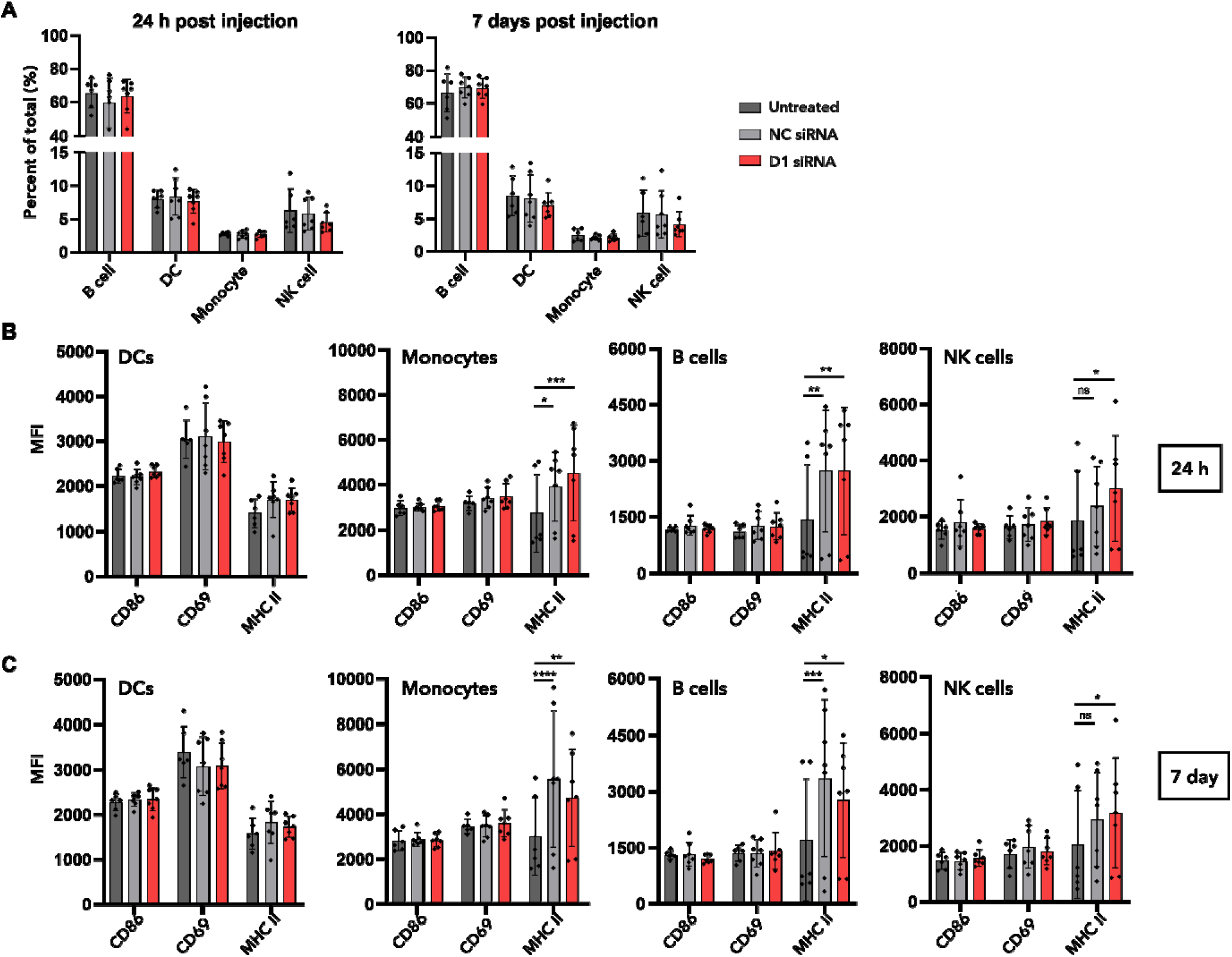
MHC class II expression on peripheral NK cells is increased in mice implanted with D1 siRNA treated cells. (**A-C**) Mice were treated as in the schematic in Figure 2A. At 24 h and 7 days post implantation, mice from each treatment group were bled, and peripheral (circulating) blood mononuclear cells were isolated. Cell type and activation status were assessed by antibody staining and analyzed by flow cytometry. (**A**) Percent of circulating B cells, dendritic cells (DC), monocytes, and NK cells at 24 h and 7 days post implantation. Percent cell type relative to all circulating mononuclear cells is reported on the *y*-axis. Plotted values represent mean ± SD for n=6 (untreated) or 7 (NC and D1 siRNA treated) mice from each treatment group. No changes in percentage of circulating blood cells were observed at either timepoint. (**B-C**) Activation status of each cell type at 24 h (B) and 7 days (C) post implantation as determined by mean fluorescence intensity (MFI, reported on the *y*-axis) of CD86, CD69, and MHC class II. Plotted values represent mean ± SD for n=6 (untreated) or 7 (NC and D1 siRNA treated) mice from each treatment group. MHC class II expression was upregulated on various cell types in mice treated with siRNA when compared to untreated cells; however, only NK cell upregulation was specific to treatment with D1 siRNA. For all, statistical analyses were performed using a one-way ANOVA with post hoc multiple comparisons test corrected using the Dunnett method (*p < 0.05, **p < 0.01, ***p < 0.001, ****p < 0.0001).

**Figure S6.**
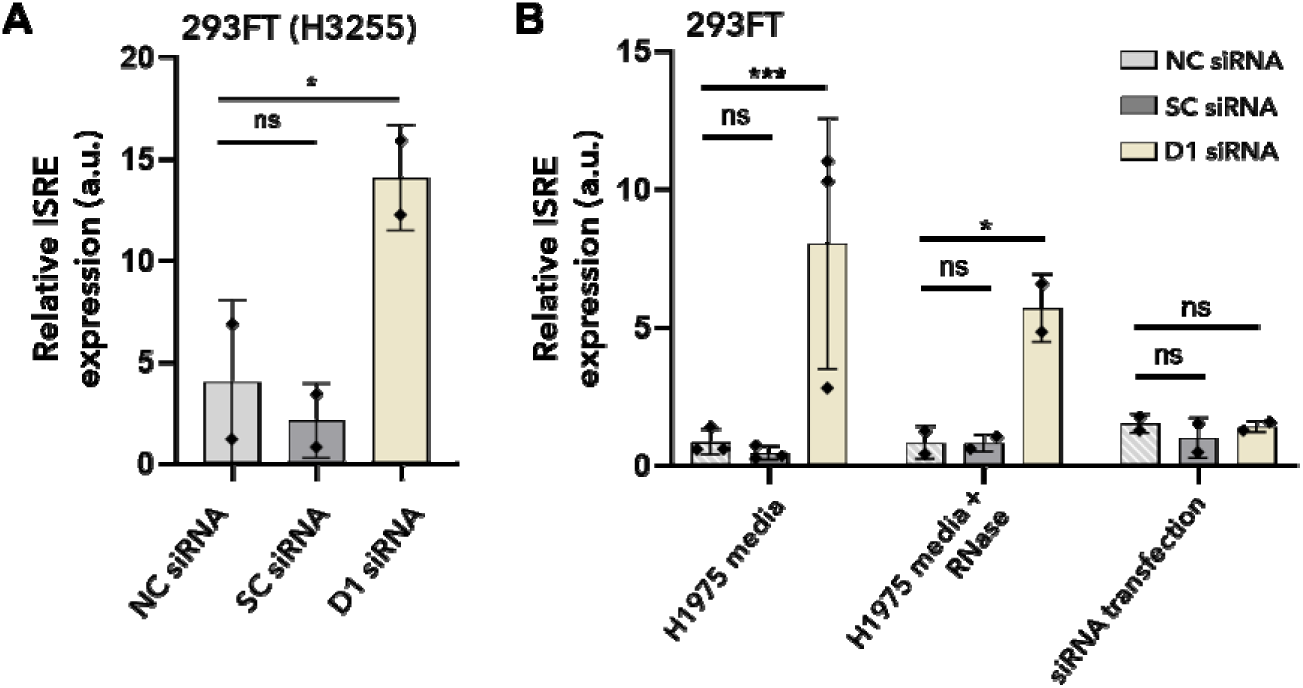
Media from DUSP11 siRNA treated cells induces ISG expression in HEK293FT. (**A**) HEK293FT cells transfected with an ISRE reporter were treated and assessed essentially as in Figure 3B, but here using media from H3255 cells treated for 48 h with 20 nM NC (light grey bar), SC (dark grey bar), or D1 (sand bar) siRNA. Cells treated with conditioned media from D1 siRNA treated cells exhibited increased ISRE expression compared to cells treated with control siRNA. (**B**) HEK293FT cells transfected with an ISRE reporter were treated and assessed as in Figure 3B, but here using media from H1975 cells transfected with 10 nM NC (light grey bar), SC (dark grey bar), or D1 (sand bar) siRNA for 48 h then treated with or without RNase (to remove any residual siRNA), or with media freshly spiked with Lipofectamine-10 nM siRNA complex. Cells treated with conditioned media from D1 siRNA treated cells exhibited increased expression of the ISRE-regulated reporter relative to control treated cells regardless of endpoint RNase exposure. In contrast, direct siRNA transfection of HEK293FT did not induce any significant changes or upregulation in reporter expression. For A-B, reporter expression was determined by luciferase substrate conversion (light production) relative to untreated cells (*y*-axis; a.u. = arbitrary units). Plotted values represent mean ± SD for n=2-3 independent experiments, each with 2-3 technical replicates. For A-B, statistical analyses were performed using a one-way ANOVA with post hoc multiple comparisons test corrected using the Dunnett method (***p < 0.001, ****p < 0.0001).

**Figure S7.**
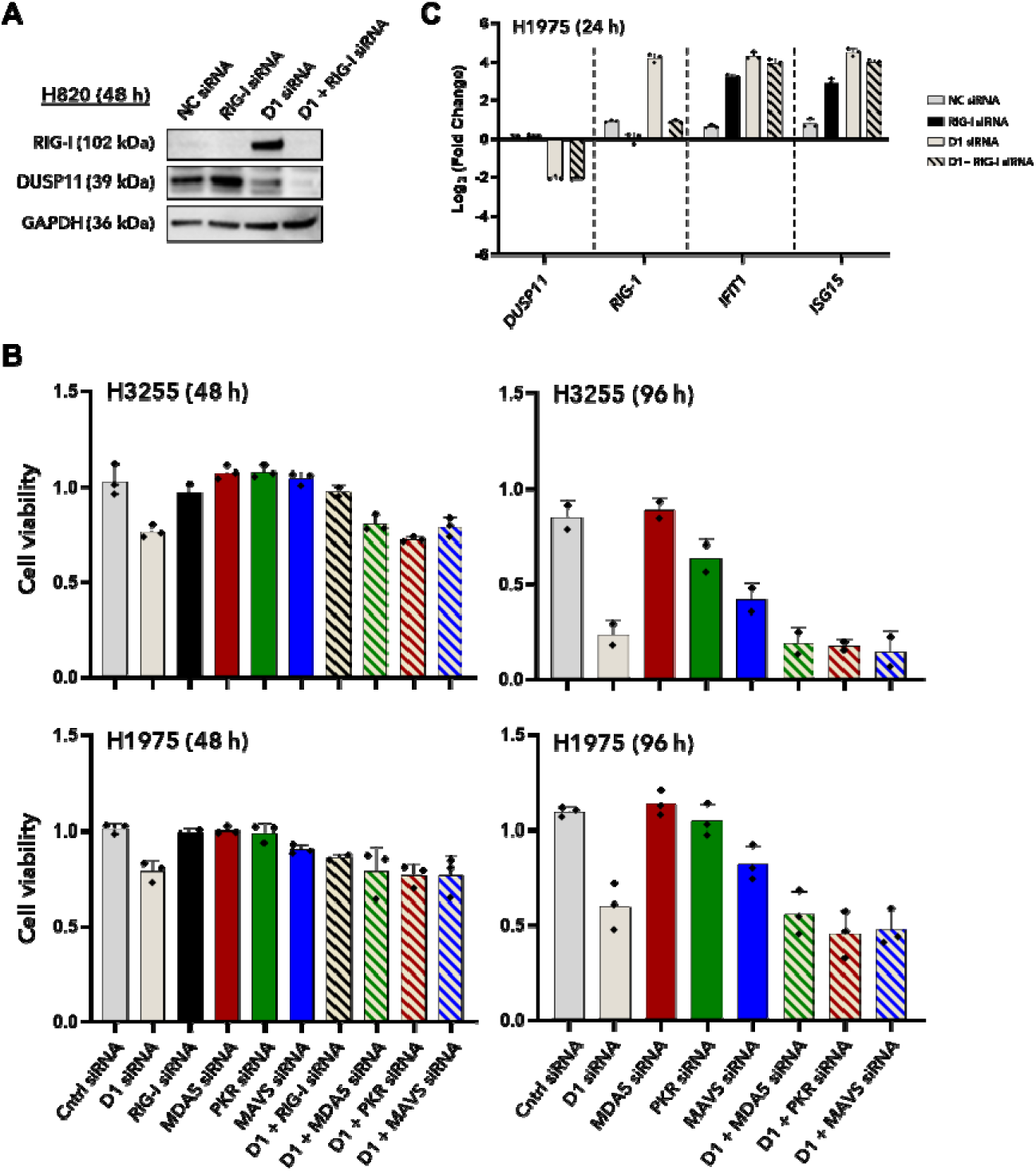
RIG-I, but not other pattern recognition receptors, is required for the innate immune response and cell viability phenotype in lung adenocarcinoma. (**A**) H820 cells were treated as in Figure 4C. In brief, cells were treated with 10 nM NC (grey bar) or D1 (sand bar) siRNA in the presence or absence of 10 nM RIG-I siRNA (black or black/sand bar) for 48 h and protein (right) expression from total cell lysate was assessed by western blot. Representative blots from n=2 independent experiments are shown. Target proteins were knocked down as expected relative to the loading control, GAPDH. (**B**) H3255 (*top*) or H1975 (*bottom*) cells were treated with 10 nM NC (grey bar) or D1 (sand bar) siRNA in the presence or absence of 10nM RIG-I siRNA (black or black/sand bar), MDA5 (red or red/sand bar), PKR (green or green/sand bar), or MAVS (blue or blue/sand bar) siRNA for 48 h (*left*) or 96 h (*right*), and cell viability was assessed by MTS assay relative to untreated cells (*y*-axis). Plotted values represent mean ± SD for n=2-3 independent experiments, each with 2-4 technical replicates. Cells co-transfected with only RIG-I siRNA (also see Figure 4A, but not MDA5 or PKR siRNA, partially recovered the cell viability phenotype. (**C**) H1975 cells were treated as in Figure 4C but mRNA was assessed by RT-qPCR at 24 h. Cells treated with RIG-I siRNA exhibited minimal RIG-I knockdown relative to controls and compared to 48 h (*see* Figure 4C).

**Figure S8.**
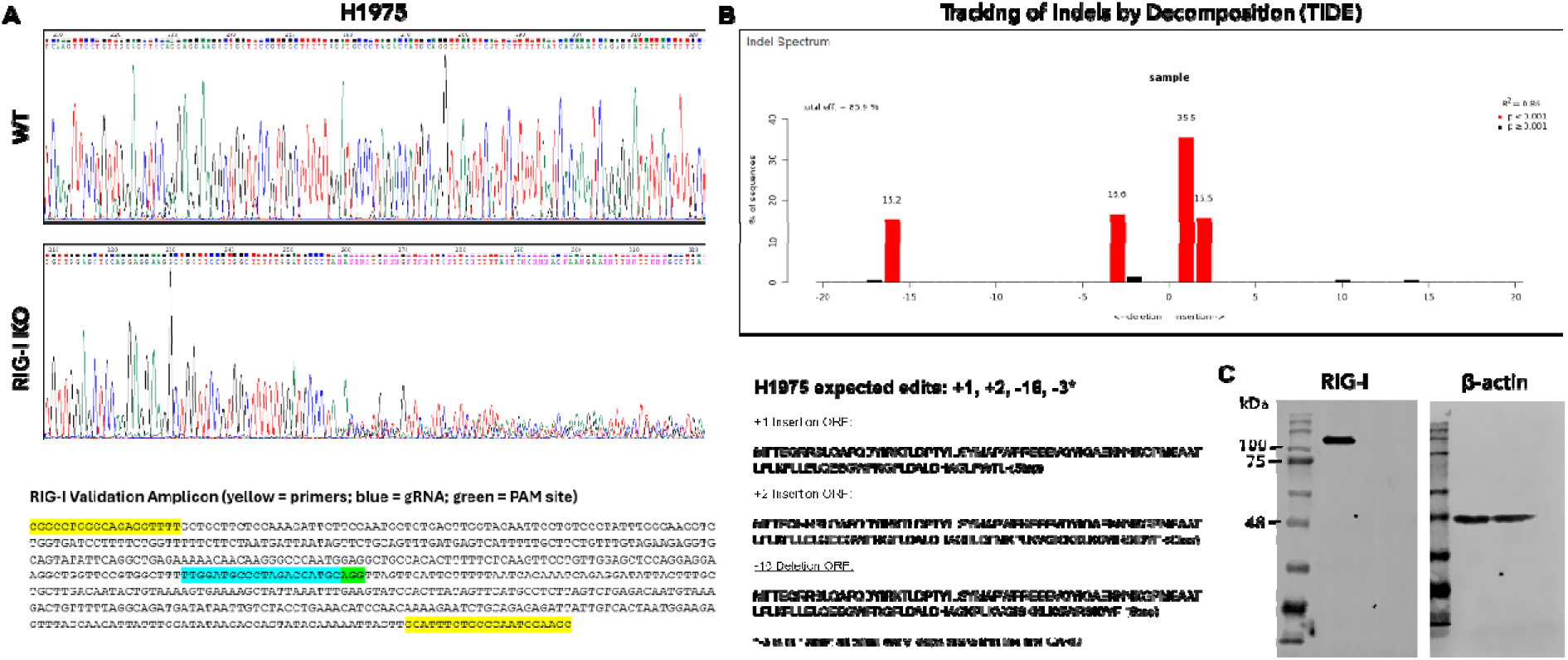
Engineering of RIG-I knockout H1975 cells. (**A**) Sanger sequencing chromatograms of H1975 WT (parental) and RIG-I KO cell lines. The RIG-I validation amplicon used for Sanger sequencing and editing confirmation is shown below with amplification primers (yellow), guide RNA (cyan), and PAM site (green) highlighted. (**B**) Tracking of Indels by Decomposition (TIDE) based on sequencing results. Expected edits and protein products are shown below. The RIG-I KO cell population has three out of frame edits (+1, +2, -16) that introduce early stops within the first caspase and activation recruitment domain (CARD). (**C**) Total cell lysates from H1975 WT (lane 1) or RIG-I KO (lane 2) were probed for RIG-I and a loading control, β-actin. A representative western blot is shown and confirmed loss of RIG-I antibody detection RIG-I KO cells as compared to WT cells.

**Figure S9.**
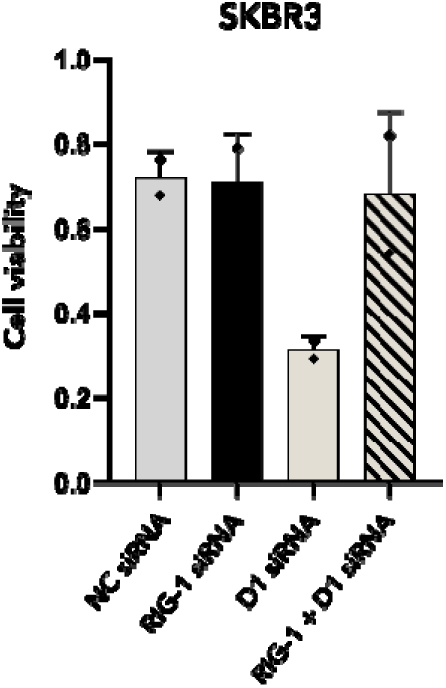
RIG-I is important for the cell viability phenotype in the breast cancer cell line SKBR3. (A) SKBR3 cells were transfected with 10 nM NC (grey bar) or D1 (sand bar) siRNA in the presence or absence of 10nM RIG-I siRNA (black or black/sand bar) for 96 h (media change with no siRNA at 48 h), and cell viability was assessed by MTS assay. Cell viability relative to untreated cells is reported on the *y*-axis and plotted values represent mean ± SD for n=2 independent experiments, each with 3 technical replicates. Cell viability, which decreased upon treatment with D1 siRNA, was recovered when cells were co-transfected with RIG-I siRNA.

**Figure S10.**
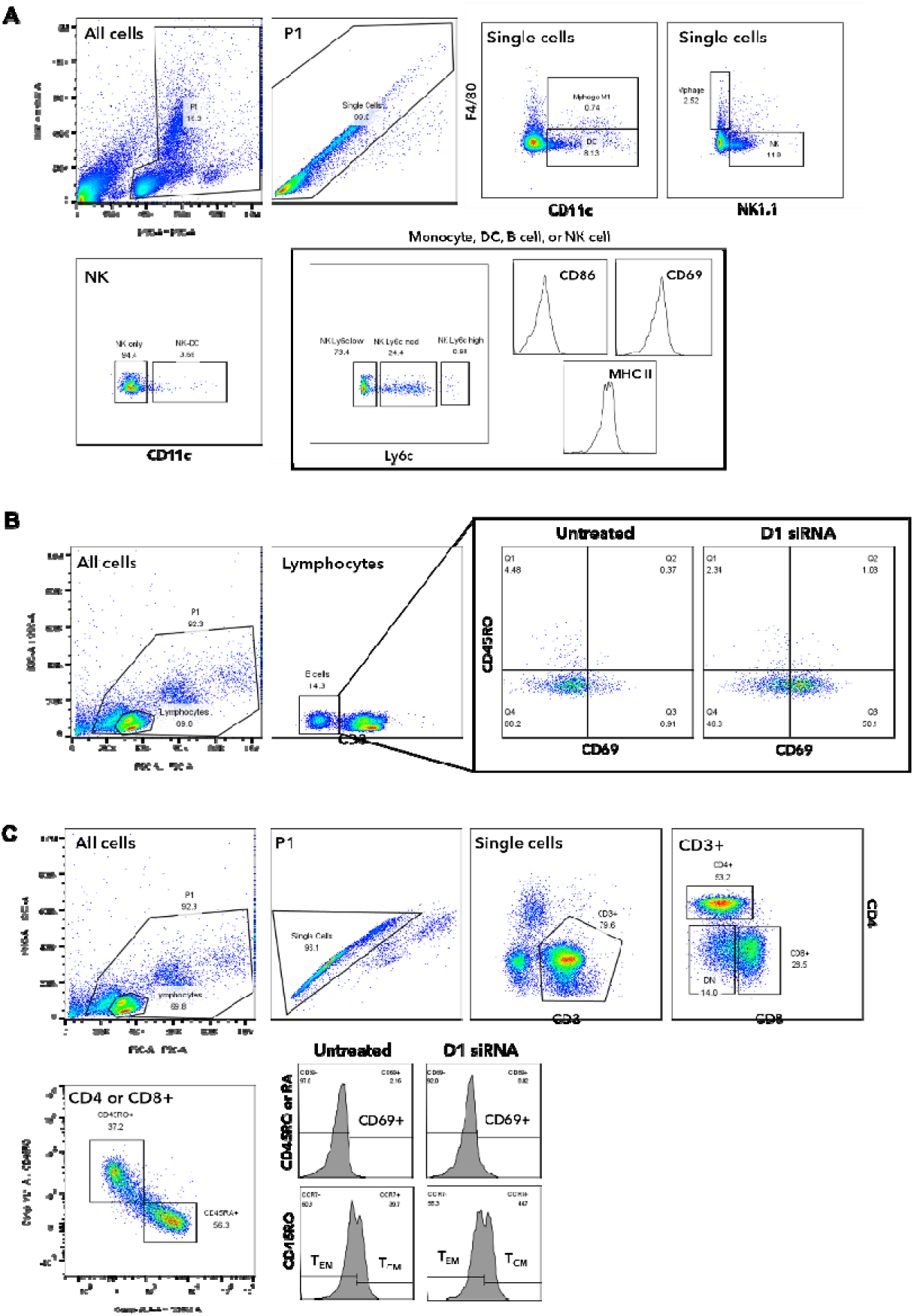
Flow cytometry gating strategies of mouse and human peripheral blood mononuclear cells. (**A**) Representative gating strategy used for BALB/c nude mouse peripheral (circulating) blood cells to remove debris (“P1” gate on ungated population) and doublets (“Single cells” gate on P1 population). Single cells were used to identify specific cell populations: dendritic cells (DC; CD11c+/F4/80-), NK cells (NK only; NK1.1+/F4/80-/CD11-), monocytes (Mphage; F4/80+/NK1.1-), and B cells (F4/80-/NK1.1-/CD11c-). Note that BALB/c nude mice are T cell deficient. Cell populations were then assessed for mean fluorescence intensity (MFI) of CD86, CD69, and MHC class II, as well as for Ly6c expression (low, mod, high). (**B-C**) Representative gating strategy used for human peripheral blood mononuclear cells to remove debris (“P1” gate on ungated populations) or non-lymphocytes (“lymphocytes” gate on ungated population) and doublets (“Single cells” gate on P1 population). Lymphocytes were used to identify B cells (CD3-), and single cells were used to identify CD4+, CD8+, or double negative (DN; CD4-/CD8-) T cells (CD3+). T cell populations were further gated for CD45RO+ (memory) or CD45RA+ (naïve). Memory T cells were further gated as either central memory (T_CM_; CCR7+) or effector memory (T_EM_; CCR7-). Cell populations were then assessed for MFI of activation marker CD69.

**Table S1.**
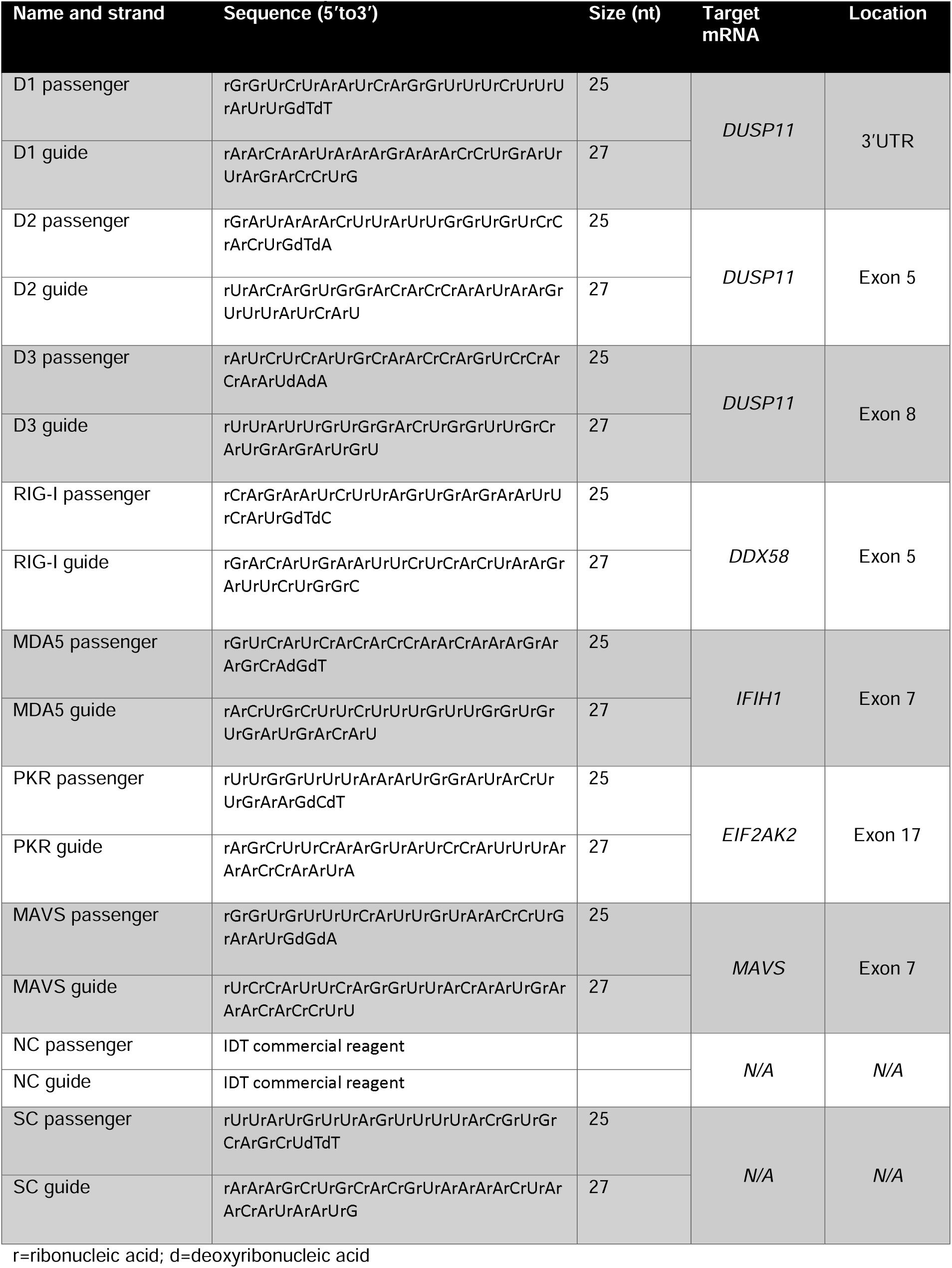
Dicer substrate siRNA sequences.

**Table S2.**
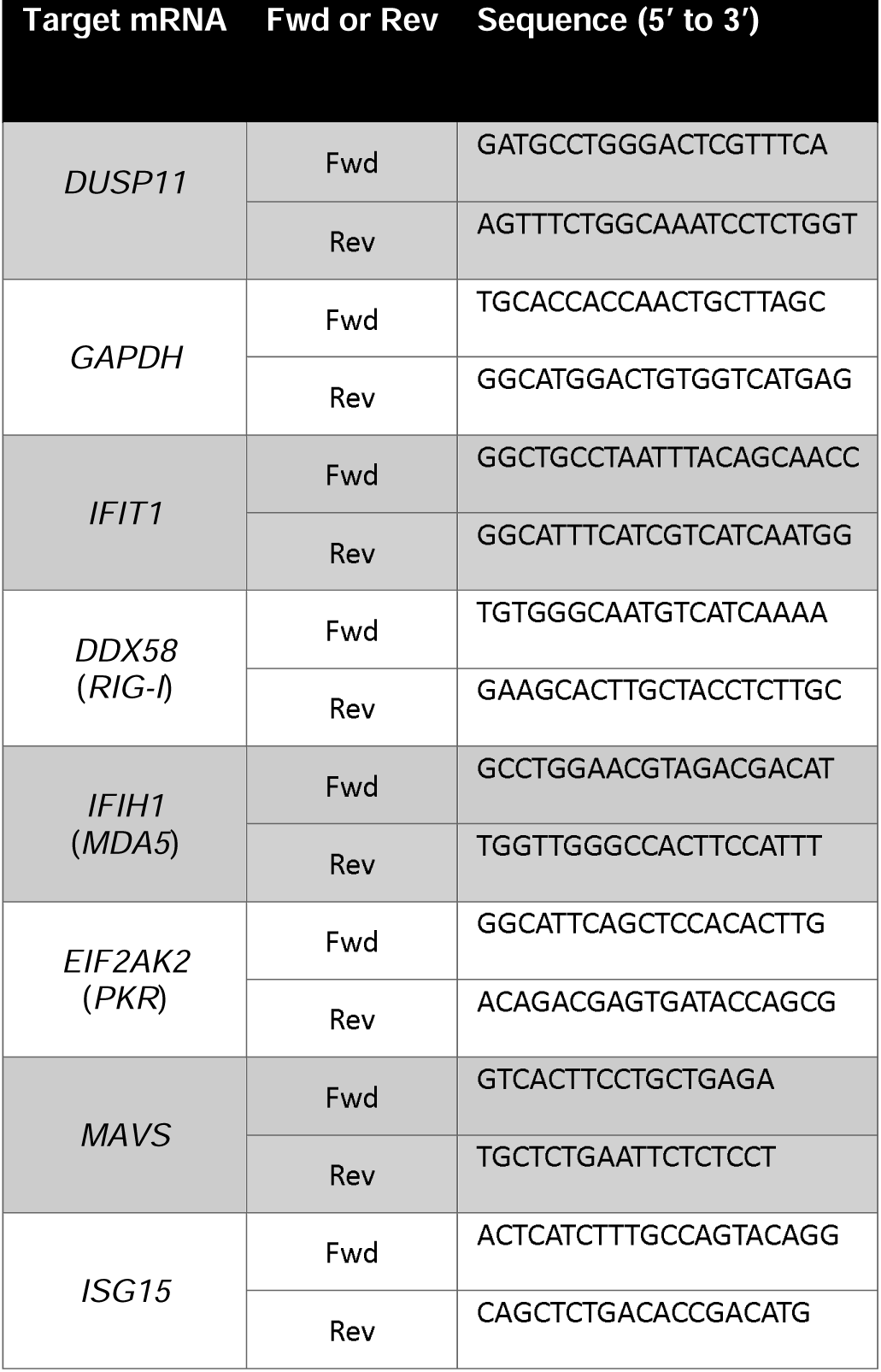
RT-qPCR primer sets.

